# Benzoxazinoid-mediated microbiome feedbacks enhance Arabidopsis growth and defence

**DOI:** 10.1101/2024.10.21.619081

**Authors:** Katja Stengele, Lea Stauber, Lisa Thoenen, Henry Janse van Rensburg, Viola D’Adda, Klaus Schlaeppi

**Affiliations:** Department of Environmental Sciences, University of Basel, 4056, Basel, Switzerland; Institute of Plant Sciences, University of Bern, 3013, Bern, Switzerland

**Keywords:** *Arabidopsis thaliana*, *Botrytis cinerea*, microbiome feedback, plant-soil feedback, root microbiota, secondary metabolites

## Abstract

- Plants modulate their surrounding microbiome via root exudates and such conditioned soil microbiomes feed back on the performance of the next generation of plants. How plants perceive altered soil microbiomes and modulate their performance in response to such microbiome feedbacks however remains largely unknown.
- As tool to condition contrasting microbiomes in soil, we made use of two maize lines, which differ in their ability to exude benzoxazinoids. Based on these differentially conditioned soil microbiomes we have established a model system with *Arabidopsis thaliana* (Arabidopsis) to investigate the mechanisms of microbiome feedbacks.
- Arabidopsis plants responding to the benzoxazinoid-conditioned soil microbiome grew better and were developmentally more advanced. Further, these plants harboured differential root bacterial communities, showed enhanced defence signatures in transcriptomes of their shoots and they were more resistant to the fungal pathogen *Botrytis cinerea*.
- Intriguingly, Arabidopsis responded with both improved growth and enhanced defence to the benzoxazinoid-conditioned soil microbiome, and we found that this simultaneous increase of growth and defence was mediated by priming of the defences. Further advancing our basic understanding how plants respond to soil microbiomes and mediate their feedbacks is particularly important for the goal to improve crops so they can benefit from their soil microbiome.

## Introduction

Plants condition the surrounding soil throughout their lifetime to optimize their immediate environment by changing the biotic and abiotic properties of the soil they grow in (van der Putten *et al*., 2013). These soil changes also persist and influence the performance of a next generation of plants, which is known as a plant-soil feedback (PSF; Bezemer *et al*., 2006; van der Putten *et al*., 2013; Mariotte *et al*., 2018). In the conditioning phase of PSFs, the growing plants alter not only the physiochemical soil properties such as soil structure, organic matter content or nutrient levels, but they also alter the specific metabolite profile (Delory *et al*., 2024), and the biotic soil properties including the soil microbiome (Semchenko *et al*., 2022; Jing *et al*., 2022). These soil alterations then “feed” back on the growth and/or defence of the next plant generation. If these feedbacks can be specifically assigned to altered soil microbiomes, which the plant perceives and responds to, we refer to them as ‘microbiome feedbacks’ (Janse van Rensburg *et al*., 2024).

The microorganisms living in soil also present the majority of the microorganisms that colonize plant roots (Spooren *et al*., 2024). These microorganisms collectively function as a microbiome, which includes bacteria, fungi, viruses and protists (Bai *et al*., 2022). The root microbiome affects plant health and performance (Bulgarelli *et al*., 2013; Trivedi *et al*., 2020), for example by regulating plant nutrient homeostasis (Salas-González *et al*., 2021), mitigating abiotic stress (Schmitz *et al*., 2022) or protecting against pathogens (Durán *et al*., 2018). To foster such beneficial functions, plants can selectively modulate the composition and activity of their root and surrounding soil microbiome through the secretion of root exudates (Sasse *et al*., 2018; Jacoby *et al*., 2021). These exudates include primary and secondary metabolites, which act as nutrient sources or specific communication cues to microbes (Sasse *et al*., 2018; Korenblum *et al*., 2022). Several specific classes of exuded metabolites have been demonstrated to structure the plant root and rhizosphere microbiota, including coumarins (Stringlis *et al*., 2018), flavones (Yu *et al*., 2021), and benzoxazinoids (Hu *et al*., 2018; Cotton *et al*., 2019; Kudjordjie *et al*., 2019).

Benzoxazinoids (BXs) are indole-derived alkaloids that function as secondary metabolites, and they are mainly produced by Poaceae, including maize, wheat and rye (Niemeyer, 2009; Niculaes *et al*., 2018). BXs are multifunctional and act for instance in the aboveground defence against generalist herbivores (Wouters *et al*., 2016; Robert & Mateo, 2022). BXs are also exuded in substantial amounts into soil, particularly by maize (Hu *et al*., 2018), where the main exuded metabolites are 2,4-dihydroxy- 7-methoxy-1,4-benzoxazin-3-one (DIMBOA) and its glucoside. DIMBOA is spontaneously converted into 6-methoxybenzoxazolin-2(3H)-one (MBOA) in soil, which is then further metabolised to 2-amino- 7-methoxyphenoxazin-3-one (AMPO) by microbes (Niemeyer, 2009). In soil, BXs have important belowground functions with their allelopathic effects on neighbouring or successional plants (Teasdale *et al*., 2012; Schandry & Becker, 2020). Typical phytotoxic compounds are the BX breakdown products 2-amino-3H-phenoxazin-3-one (APO) and AMPO, which have been shown to inhibit growth of Arabidopsis *in vitro* (Venturelli *et al*., 2015).

BXs also have important belowground functions on the microbiota in PSFs. The root and rhizosphere microbiotas are selectively shaped by the exuded BXs (Hu *et al*., 2018; Cotton *et al*., 2019; Kudjordjie *et al*., 2019), which leaves a legacy in soil for a next plant generation. This BX-dependent soil conditioning was found to affect growth and defence of a next generation of maize (Hu *et al*., 2018) or wheat (Cadot *et al*., 2021). Sterilisation and microbiota complementation experiments demonstrated that these feedbacks were mediated by the microbes, and supplementation of MBOA to non-BX exuding maize plants established that soil conditioning was BX-dependent (Hu *et al*., 2018). Hereafter, we refer to these PSFs as ‘BX-feedbacks’. BX-feedbacks are studied by comparing soil conditioned with wild-type maize plants that exude BXs, called ‘BX_plus_’ soil, to the control ‘BX_minus_’ soil that is conditioned by the BX-defective mutant line *bx1* (Maag *et al*., 2016). While the phenology (Hu *et al*., 2018; Cadot *et al*., 2021) and the agronomic relevance (Gfeller *et al*., 2023b,a) of BX-feedbacks have been well described, the mechanistic understanding of how plants perceive the differential soils and how they alter their own performance remains largely unknown. In more general terms, how plants perceive complex soil microbiomes and how they mediate microbiome feedbacks, is currently understudied (Janse van Rensburg *et al*., 2024).

In search of a model system to investigate microbiome feedbacks and to facilitate mechanistic studies, we explored BX-feedbacks with the model plant *Arabidopsis thaliana* Col-0 (henceforth Arabidopsis), which we grew on BX_plus_ and BX_minus_ soils. Arabidopsis robustly responded with increased shoot growth on BX_plus_ soil. Through sterilisation, we corroborated that the soil microbiota is required for this growth feedback. BX-conditioning also lead to different bacterial communities on Arabidopsis roots. A transcriptome analysis revealed upregulation of defences in shoots, which coincided with smaller lesions of the pathogen *Botrytis cinerea* and increased *PR1* expression after treatment with salicylic acid when plants were grown on BX_plus_ soil. Thus, we uncovered that the BX_plus_ soil microbiota primed the Arabidopsis’s defences, providing an explanation for the better resistance to Botrytis alongside the simultaneous improved growth feedback.

## Materials and Methods

### Plant material

For the soil conditionings, we used the maize (*Zea mays* L.) variety B73 and the benzoxazinoid-deficient line *bx1* in B73 background (Maag *et al*., 2016). For all feedback and in vitro experiments, we used the *Arabidopsis thaliana* accession Columbia-0 (Col-0).

### Soil conditioning

We collected soil from three neighbouring fields managed by Agroscope in Changins (Nyon, Switzerland) between 2019 and 2022 to condition with maize. Soils were sieved (1 cm mesh size), homogenized, and stored at 4°C before conditioning. For each conditioning event, which yields one ‘soil batch’, we grew wild-type B73 and *bx1*(B73) maize for 12 weeks in individual pots. For details on the set up of each conditioning experiment see **Table S1**. Plants were watered three times a week and supplemented once a week with nutrient solution. After twelve weeks, the roots were removed from the pots and the soils were collected and pooled by maize genotype, resulting in BX_plus_ soil (i.e., conditioned with B73 wild-type maize) and BX_minus_ control soil (i.e., conditioned with *bx1*(B73) maize). Soils were stored at 4°C until further use (see **Table S2** for soil storage duration of each experiment).

### Feedback experiments in soil

As a general procedure to perform feedback experiments with Arabidopsis using soil, the conditioned soils were sieved (5 mm mesh size), amended with 20% (v/v) autoclaved sand and filled to the same weight into small pots. Arabidopsis seeds were stratified for 2-3 days either directly on the pots or suspended in 0.2% sterile agar before sowing. Several seeds were distributed per pot, and after 1-2 weeks, the healthiest looking seedling was kept and the others were removed from the pot. Plants were grown in growth cabinets (CLF Plant Climatics, Wertingen, Germany) at 60% relative humidity, 10 h day at 21°C and light intensities between 100 and 200 µmol m^-2^ s^-1^, and 14 h night at 18°C. Short-day conditions were chosen because slower plant development would allow longer exposure of roots to the soil microbiome. Pots were watered with tap water three times per week based on weight, and the pot position was changed once per week. Plants were fertilised by watering with 1/3 half strength Hoagland solution (Cardoso *et al*., 2014) diluted in tap water. Plants were grown for six to seven weeks before harvesting. Here we described the general procedure applying to all experiments, while we summarised the different setups of the experiments (Exp. I to Exp. XII) in detail in **Table S2**, and we explain additional details of particular experiments in the **Supplementary Methods**. For all experiments, rosette area was determined by either analysing scans of cut rosettes with ImageJ v1.5 (Schneider *et al*., 2012), or by analysing photographs of rosettes from living plants with the software ARADEEPOPSIS (Hüther *et al*., 2020).

### In vitro experiments

We have performed three in vitro experiments, two on agar plates and one using a semi-hydroponic growth system (McLaughlin *et al*., 2023), to test the effects of MBOA on Arabidopsis growth and resistance to Botrytis. The three experiments were conducted using the same Percival growth chambers and the same growth settings as for the feedback experiments in soil. The specific experimental details and measurements are documented in the **Supplementary Methods**.

### Infections with *Botrytis cinerea*

For infections, the *Botrytis cinerea* strain B05.10 was grown in the dark at 18°C on potato dextrose agar (Merck Millipore, Burlington, USA) for two to three weeks. Plants were grown as described under *Feedback experiments in soil*. At five to six weeks, three mature source leaves per plant were infected with Botrytis spores that were harvested from plates with sterile water + 0.0001% Tween-20 (Biorad, Basel, Switzerland), filtered through glass wool, and adjusted to 1x10^5^ spores per mL in ½ strength potato dextrose broth (Formedium, Norfolk, United Kingdom). For infection, a 5 µL drop was placed onto the middle of the leaf towards the edge. The plant trays were covered with a water-sprayed transparent lid that was taped to the tray to ensure high humidity. For the infection, plants were transferred to a growth chamber (Sanyo, Moriguchi, Japan) at the same growth conditions as previously, but equipped with fluorescent bulbs of ∼ 100 µmol m^-2^ s^-1^ light intensity. Experiment XI was incubated in a Sanyo chamber equipped with LEDs under low light conditions (∼ 50 µmol m^-2^ s^-1^) for the infection. Photographs from the infected leaves were taken from the top at a fixed distance three days after the infection. The area was quantified using the Fiji distribution of Image J (Schindelin *et al*., 2012) by selecting the area of the lesions with the Polygon tool and calculating the area of the lesions.

### Defence priming of *PR1*

The shoots of six-week-old plants (Experiment VII) were dipped in a 1mM SA solution containing 0.015% Silwet, L-77, or mock treated by dipping in tap water containing 0.015% Silwet L-77. After 30 minutes, 4, 6 and 24 hours, shoots were collected, snap frozen in liquid nitrogen and stored at −80°C. The whole shoots were ground under liquid nitrogen, and RNA was extracted with the RNeasy plant kit (Qiagen, Hilden, Germany). cDNA was then synthesized with the GoScript cDNA Synthesis kit (Promega), and qPCR reactions with the KAPA SYBR® FAST qPCR Master Mix (Kapa biosystems, Wilmington, USA) were set up for *PR1* and the reference gene *PP2A* for each sample in duplicate. Per 15 µL reaction, 0.3 µL of the *PP2A* primers (Rodriguez *et al*., 2010) and the *PR1* primers (Pieterse *et al*., 1998) were used at 10 µM each. Quantitative real-time PCR (qPCR) was performed on a LightCycler® 480 (Roche, Basel, Switzerland) following the recommended cycling parameters from KAPA with an annealing temperature of 60°C for 25 seconds followed by fluorescence acquisition for 2 seconds at 72°C. The data was then processed with the LinRegPCR software (Ruijter *et al*., 2009) and further analysed in R v4.3.0 (R Core Team, 2017).

### Determination of benzoxazinoid concentrations in soils

Before the first Arabidopsis experiments, soil samples of soil batches 1, 2 and 4 were stored at −80 °C, to analyse the BX concentrations following the sample processing and analysis described in Gfeller et al. (2023b). Briefly, BXs were extracted with acidified Methanol (70% Methanol, 0.1% formic acid), and analysed in an Acquity UHPLC system coupled to a QTOF mass spectrometer without further concentration of samples. Pure BX compounds as described in Gfeller et al. (2023b) were used to determine absolute quantities via calibration curves.

### Microbiota profiling

For microbiota profiling, we analysed the bacterial and fungal communities of roots grown in native and sterilised soils along with native soil samples (Experiment III). Sampling, DNA extraction, PCR, library preparation, sequencing and the bioinformatic analysis are detailed in the **Supplementary methods**. The microbiota profiles were analysed in R v4.3.0 (R Core Team, 2017). We first removed reads assigned to Eukarya, Archaea, Cyanobacteria or Mitochondria from the bacterial amplicon sequence variant (ASV) count table. Due to low read numbers, we removed 11 samples from the ITS library (<900 reads). We normalised both libraries with total sum scaling and did not rarefy the libraries, as the number of reads did not differ between sample groups (Weiss *et al*., 2015). We ran data ordinations based on Bray-Curtis dissimilarity matrices (Bray & Curtis, 1957) using the package phyloseq v1.44.0 (McMurdie & Holmes, 2013) and visualized them with unconstrained Principal Coordinate Analysis (PCoA). We performed Permutational Multivariate Analysis of Variance (PERMANOVA) with 99’999 permutations with the adonis function from the vegan package to partition and quantify effects of our experimental factors. To determine differentially abundant ASVs, we used four different identification methods, implemented in the packages ALDEx2 v1.34.0 (Fernandes *et al*., 2013, 2014), Maaslin2 v1.14.1 (Mallick *et al*., 2021), metagenomeSeq v1.43.0 (Paulson *et al*., 2013), and ANCOMBC v2.2.2 (Lin & Peddada, 2020). We defined an ASV to be differently abundant if it was detected (adjusted *p*-value < 0.05 using the Benjamini–Hochberg method) by at least three of the four methods. Data and code are made public (see below).

### Transcriptome analysis

Arabidopsis root and shoot sample material were collected for the RNAseq analysis of seven-week-old plants from Experiment V.II. Whole shoots were directly frozen in liquid nitrogen, and the whole roots were collected and removed from all loosely attached soil before freezing. The RNA was extracted with the NucleoSpin RNA kit (Macherey-Nagel, Düren, Germany) and quantified via Fluorometry. Next, RNA quality was checked with TapeStation, and prepared as a library with 200 ng total RNA input using polyA enrichment with the TruSeq Stranded mRNA kit (Illumina, SanDiego, USA). The library was sequenced with a 2x 51 bp paired-end protocol with Illumina NovaSeq6000 (Illumina, San Diego, USA) at the Genomics Facility Basel (https://bsse.ethz.ch/genomicsbasel).

After quality control, reads were aligned to the Arabidopsis thaliana TAIR10 reference genome and read count was performed using STAR 2.7.9a (Dobin *et al*., 2013). All raw data processing was performed at sciCORE (http://scicore.unibas.ch/) scientific computing core facility at University of Basel. Downstream analyses were performed with R 4.3.0 using DESeq2 v1.42.1 (Love *et al*., 2014) for differential gene expression analysis. We considered genes to be differentially expressed if the adjusted *p*-value (*p*_adj_) < 0.05 and log_2_FoldChange > ±1. Co-expressed genes were identified using the package coseq v1.24.0 (Rau & Maugis-Rabusseau, 2018). Models were rerun 20 times to determine the optimal number of clusters. Genes with conditional cluster membership probabilities ≥0.95 and *p*_adj_ < 0.01 were then used for cluster-wise gene ontology (GO) enrichment analysis using hypergeometric testing, as implemented in the package clusterProfiler v4.10.0 (Wu *et al*., 2021). Data and code are made public (see below).

### General statistical analyses

All statistical analyses were executed in R v4.3.0 (R Core Team, 2017) within Rstudio v2023 (Posit team, 2023). Rosette area or biomass data were statistically compared between two groups using two-sided t-tests, and correlations were assessed with Pearson correlation tests. Analysis of variance test was used to compare more than one group. Data and code are made public (see below).

## Results

### Benzoxazinoid-dependent soil conditioning affects Arabidopsis growth

In search of a model system to investigate microbiome feedbacks, we tested the *Arabidopsis thaliana* accession Col-0 (hereafter, Arabidopsis) and asked if BX-conditioned soils would lead to differences in growth. We exposed Arabidopsis plants to contrasting soil microbiomes by growing them individually in pots either containing control ‘BX_minus_ soil’, where no BXs were detected, or BX- conditioned ‘BX_plus_ soil’, where MBOA and AMPO had accumulated (**Fig. S1a**). We observed that plants developed larger rosettes when grown on BX_plus_ compared to the control BX_minus_ soil (**Fig. 1a**). This observation was confirmed with the shoot fresh and dry weight and was also reflected using image-based quantification of the rosette area (**Fig. 1b**), which correlated strongly with shoot biomass (**Fig. S1b**) and was therefore, employed for the rest of the study. The positive BX-feedback on growth was reproducible with varying setups conducted with independently conditioned soil batches (**Fig. S1c**, **Table S2**) and was increasing over time (**Fig. S1d**). Furthermore, we noticed a feedback on development with increasing number of leaves at a late growth stage (**Fig. S1e**), and also recorded larger rosettes (**Fig. S1f**) and an earlier transition to flowering when plants were grown under long-day conditions on BX_plus_ soil (**Fig. 1c**). This coincided with higher seed yield on BX_plus_ soil (**Fig. 1d**; also quantified by weight, **Fig. S1g**), while the average seed weight did not differ between the two soils (**Fig. 1d**). We also investigated if BX-feedbacks affected root growth in two experiments, and found a trend to higher root biomass on BX_plus_ soil (**Fig. S1h**), which correlated significantly and positively with rosette area (**Fig. S1i**). Taken together, Arabidopsis Col-0 responded with increased growth and faster development on BX_plus_ compared to BX_minus_ soil.

**Figure 1.**
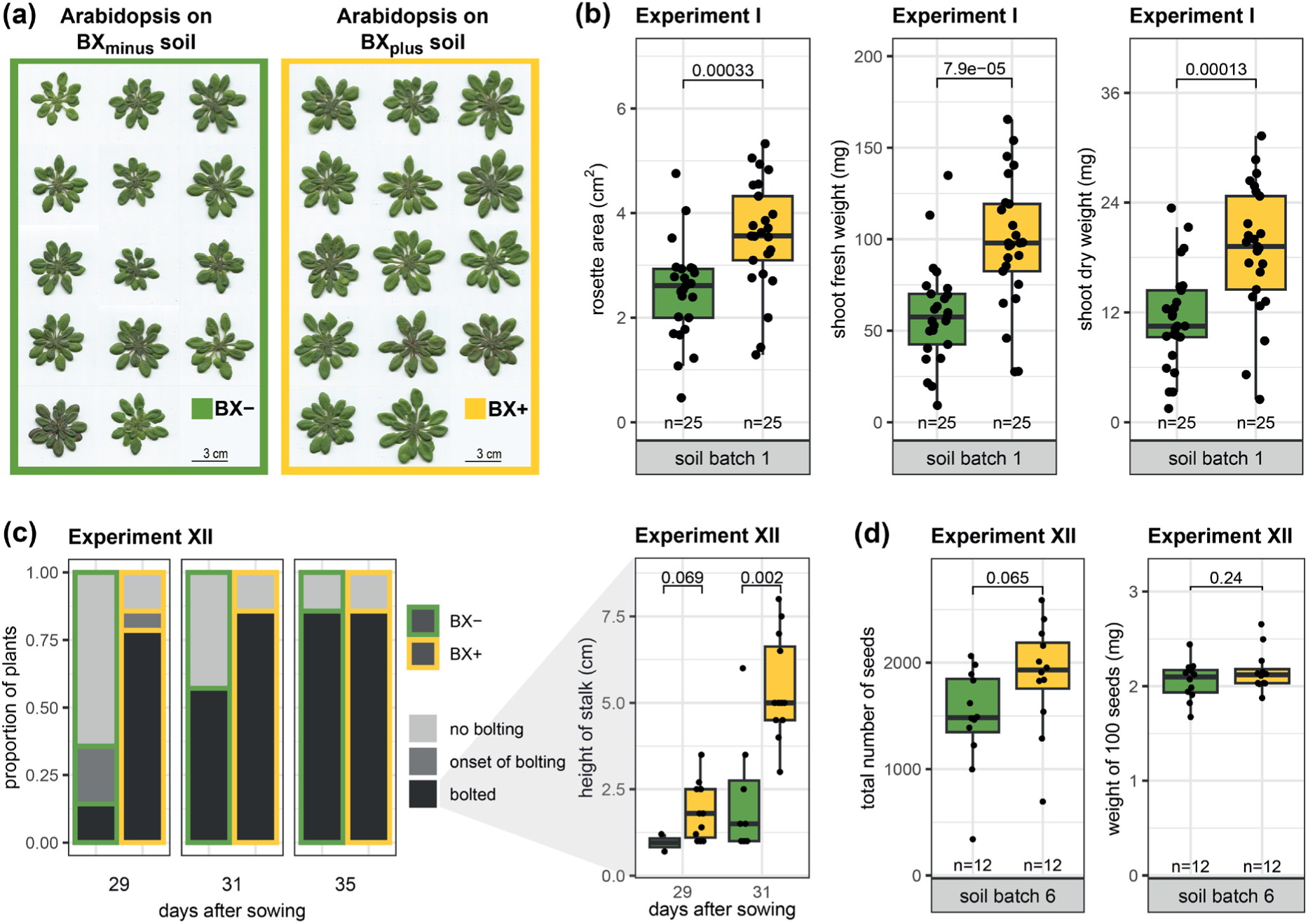
Arabidopsis growth and development is improved on BXplus compared to BXminus soil. **(a)** Harvested and scanned shoots of *Arabidopsis thaliana* Col-0 plants grown on control BXminus or BXplus soil. Photographs are from Experiment IV (see Fig. S1c). **(b)** Shoot growth measurements of Arabidopsis grown for seven weeks on BXplus and BXminus soils. From left to right, the rosette area, shoot fresh weight and shoot dry weight are shown. The boxplots further indicate the *p*-values of two-sided student’s t-tests, the replicate numbers and the soil batch number of the soil conditioning event. **(c)** Feedbacks on flowering were recorded by scoring the fractions of plants that were at the onset of bolting and bolted plants (left) and the heights of the developped flower stalks of bolted plants (right). Plant bolting was recorded at three time points (indicated at the bottom by the number of days after sowing). The height of the flower stalks of bolted plants was recorded at two time points, and the *p*-values of two-sided student’s t-tests area shown. **(d)** Measurements of seeds harvested from Arabidopsis plants that were grown on BXminus and BXplus soils. The total number of seeds of single plants were counted (left) and the weight of 100 seeds (right; calculated based on the total weight of all seeds of a plant, Fig. S1g) are shown. The boxplots further indicate the *p*-values of two-sided student’s t-tests, the replicate numbers and the soil batch number of the soil conditioning event.

### The soil microbiota mediates Arabidopsis growth differences

With maize we previously found that differential soil microbiotas were driving the BX-feedbacks based on a sterilisation experiment (Hu *et al*., 2018). To test this also with Arabidopsis, we sterilised BX_minus_ and BX_plus_ soils using X-radiation. Compared to the native soils, Arabidopsis grew overall larger on the sterilised soils (**Fig. 2a**), as sterilisation can enhance organic matter content via lysis of microbial cells (Berns *et al*., 2008). The better growth on native BX_plus_ soil was no longer seen on sterilised soil, confirming that soil biotic factors drive the BX-feedbacks. In contrast to native soils, Arabidopsis plants grew smaller on sterilised BX_plus_ compared to sterilised BX_minus_ soil. This suggested that Arabidopsis growth was reduced by a factor contained in sterilised BX_plus_ soil. The primary candidates for this factor are the BXs, as both MBOA and AMPO levels were not affected by the sterilisation (**Fig. S1a**). AMPO has been previously demonstrated for its allelopathic properties against Arabidopsis (Venturelli *et al*., 2015), thus we have also tested Arabidopsis’ sensitivity to MBOA. We did this first in sterile settings, as it would be initially the case in sterilised soil.

**Figure 2.**
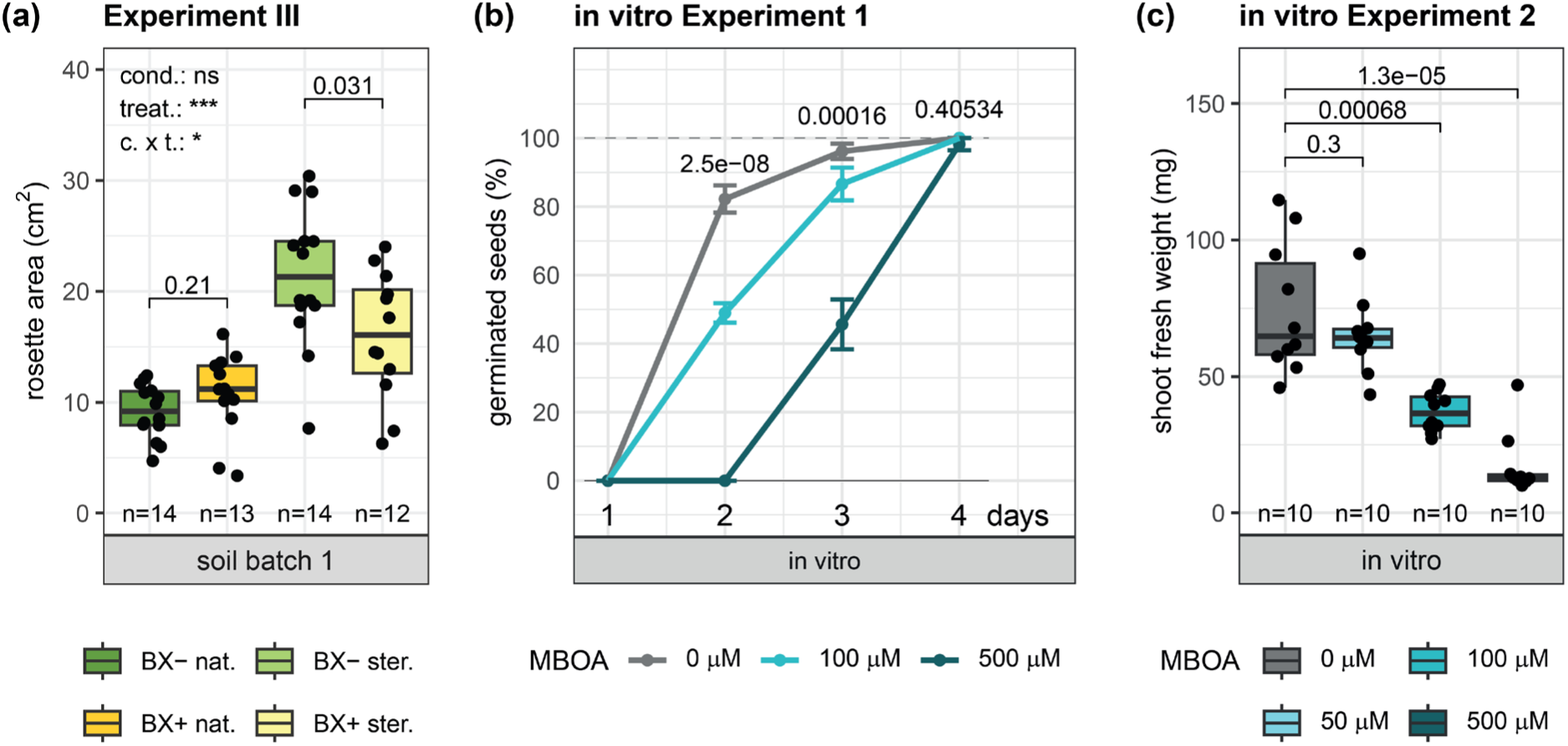
Arabidopsis no longer displays a positive growth feedback on sterilised soil and is sensitive to MBOA in vitro. **(a)** Rosette area of Arabidopsis plants grown on native (‘nat.’) and sterilised (‘ster.’) BXminus and BXplus soils. The used soil batch, *p*-values of two-sided student’s t-tests and replicate numbers are indicated with the boxplots. The significance levels of the ANOVA model testing the effect of conditioning (‘cond.’) and sterilisation treatment (‘treat.’) on rosette area are depicted in the left top corner of the graph. **(b)** Percent of germinated seeds on ½ MS plates supplemented with DMSO (‘0 µM’) or increasing amounts of MBOA (dissolved in DMSO), recorded at day 1 to 4 after sowing. The *p*-values on top of each time point represent the ANOVA statistics testing for differences between the groups. **(c)** Shoot fresh weight of Arabidopsis plants grown on ½ MS plates supplemented with DMSO (‘0 µM’) or increasing amounts of MBOA dissolved in DMSO. One data point represents the total shoot fresh weight from five plants grown on one plate. *P*-values of two-sided student’s t-tests for the respective comparisons are reported, and replicate numbers are indicated with the boxplots.

In agar plates containing MBOA, seed germination was delayed (**Fig. 2b**) and shoot biomass of three-week-old plants was reduced starting at 100 µM MBOA (**Fig. 2c**). Roots were less sensitive, as length (**Fig. S2a**) and biomass (**Fig. S2b**) were only affected at 500 µM MBOA. We then tested Arabidopsis sensitivity to MBOA in non-sterile settings. Applying MBOA to unconditioned native soil, we only found a slight reduction in shoot growth at 500 µM MBOA (**Fig. S2c**).

Thus, although Arabidopsis was sensitive to MBOA in vitro, only minor allelopathic effects and at higher MBOA concentrations were observed for Arabidopsis grown in natural soil. To conclude the sterilisation experiment (**Fig. 2a**), the biotic soil properties shaped by BX-conditioning were needed for the positive growth of Arabidopsis, highlighting that Arabidopsis BX-feedbacks represent *microbiome feedbacks*.

### Soil and Arabidopsis root microbiota compositions differ by BX-conditioning

Our previous work had also shown that differential microbiotas assembled on roots of maize plants when grown on BX_minus_ or BX_plus_ soils (Hu *et al*., 2018). To test for this with Arabidopsis, we profiled bacterial and fungal communities on roots and soil collected from the sterilisation experiment (Experiment III). We have also profiled the root microbiotas of plants grown in the initially sterilised soils, as the roots will become re-colonized by microbes during the course of the experiment in the non- sterile growth chamber.

First, we examined the communities for compositional differences between sample groups. For bacteria, ordination analysis revealed a clear separation of the communities between soil and between the two groups of roots with strongly differing communities from plants grown in native or initially sterilised soils (**Fig. 3a**). For the fungi, soil and root communities were also separated, but the separation between plants from native or sterilised soil was less marked as for the bacteria (**Fig. 3b**). Next, we quantified the effect sizes of sample groups and of the soil conditioning based on the *R^2^* values of Permutational Multivariate Analysis of Variance (PERMANOVA). Because of the overly large variation among the different sample types (bacteria 23.4%; fungi 17.8%) relative to the conditioning effects (bacteria 1.5%; fungi 1.9%; **Supplementary Dataset S1**), we tested significance of the effect sizes due to conditioning within each sample group separately. We found significantly different bacterial communities between BX_minus_ and BX_plus_ in all sample groups. BX-conditioning explained 12.2% of the bacterial community variation in soil, 7.3% on roots from native soils and 6.2% on roots from sterilised soils (**Fig. 3c**). The fungal communities only differed on roots from sterilised soil, where the BX-conditioning explained 11.7% of the fungal community variation (**Fig. 3d**). Together with the soil BX analysis (**Fig. S1a**), the microbiota profiling points to the conclusion that BXs in soil cause a differentiation of the soil and root microbiotas. This further suggests that these differential microbiotas – particularly the bacteria – may drive the differential growth of Arabidopsis.

**Figure 3.**
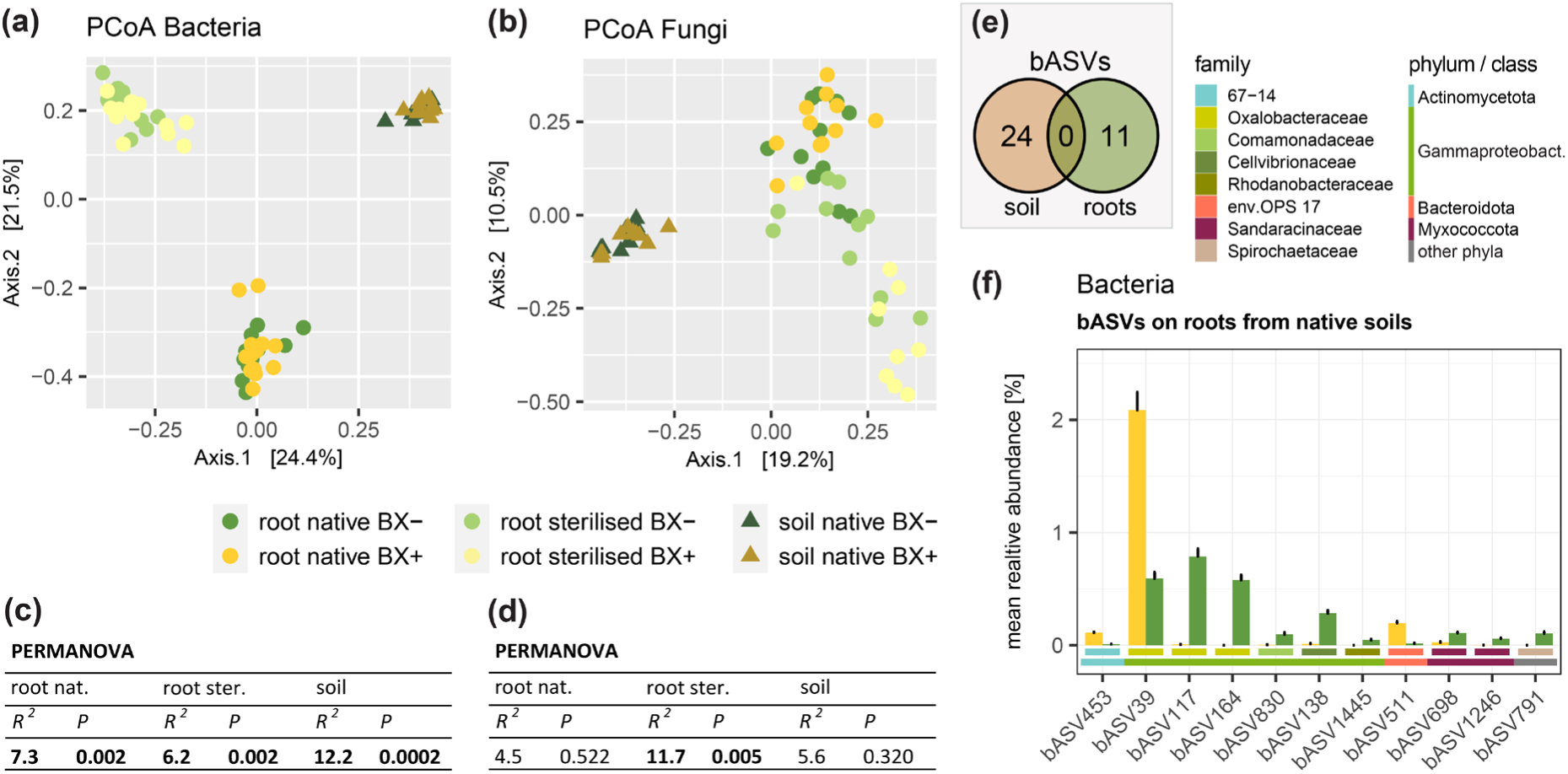
The Microbiota of soil and Arabidopsis roots differs by BX-conditioning. Principle Coordinate Analysis (PCoA) of **(a)** bacterial and **(b)** fungal community profiles from BXminus and BXplus soils and Arabidopsis roots grown in native or initially sterilised BXminus and BXplus soils. Replicate numbers are n = 10 to 14 for bacterial and n = 9 to 11 for fungal samples. PERMANOVA tables on Bray-Curtis dissimilarities of **(c)** bacterial and **(d)** fungal community profiles between the three sample groups with *R^2^* and *p*-values displayed for the conditioning variable. Statistically significant differences are shown in bold. **(e)** Overlap of bacterial ASVs (bASVs) that differed in abundance due to conditioning between native soil and roots from native soils. **(f)** Abundance and taxonomy of differentially abundant bASVs on Arabidopsis roots from native BXminus and BXplus soils. Error bars represent the standard error of the mean. The colored bars below the bargraphs indicate the bacterial family and phylum/class information (for the Pseudomonadota phylum, the class information is given, i.e. Gammaproteobacteria). The reported bASVs were identified as differentially abundant by at least three of four statistical tools used (see methods).

Next, we performed an in-depth microbiota analysis, which is documented in the **Supplementary Dataset S1**. In essence, with robust statistic support (adjusted *p*-value < 0.05) from at least three of the four utilized statistical tools, we detected in soil 24 bacterial sequences (named *b*acterial *A*mplicon *S*equence *V*ariant*s*, bASVs) whose abundances were dependent on BX-conditioning (**Fig. 3e**, **Table S3**). BX_plus_ soil was mainly enriched with 7 abundant (> 0.1% relative abundance) Sphingomonadaceae, two Gammaproteobacteria, a Chloroflexota and four Acidobacteriota bASVs, while bASVs assigned to Actinomycetota, Gammaproteobacteria and other taxonomic groups were depleted (**Fig. S3**). In roots, other 11 bASVs differed between the microbiotas of plants grown in native BX_minus_ vs. BX_plus_ soil (**Fig. 3e**, **Table S4**). Most apparent was the strong enrichment of the abundant bASV39 (genus Massilia, family Oxalobacteraceae) on roots of plants grown in BX_plus_ soil, while other ASVs of this family or of the same class were depleted (**Fig. 3f**). To summarize, consistent with maize, Arabidopsis had different bacterial communities on the roots when grown on the two soils.

### Defence-related shoot transcriptome in plants grown on BX_plus_ soil

To obtain insights into how Arabidopsis achieves better growth on BX_plus_ soil, we performed a transcriptome analysis of shoot and root samples from plants grown for seven weeks on BX_minus_ and BX_plus_ soils (**Fig. S1c**, Experiment V.II). The detailed RNAseq analysis is documented in the **Supplementary Dataset S2**. Comparing plants grown on control BX_minus_ versus BX_plus_ soil, we found 1’959 and 1’839 differentially expressed genes in shoots (**Table S5**) and roots (**Table S6**), respectively (Criteria: adjusted *p*-value, *p*_adj_ < 0.05; log_2_ fold change, log_2_FC > ±1). The subsequent co-expression and gene ontology (GO) enrichment analysis revealed that soil conditioning affected many and diverse biological processes both in shoots (**Fig. S4a**, **Table S7**) and roots (**Fig. S4b**, **Table S8**). Because many enriched GO terms were supported by rather few genes, we focused on the top two in each co-expression cluster.

In shoots, 534 genes were up- and 1’425 downregulated on BX_plus_ compared to the control BX_minus_ soil (**Fig. 4a**, **Table S5**). The main upregulated processes involved systemic acquired resistance (SAR), leaf senescence and chlorophyll and protein catabolism as well as response to abscisic acid (**Fig. 4b**). While **Tables S7** and **S8** list all genes and their links to GO term numbers, we report the statistics of exemplary genes for the mentioned GO terms in **Table S9**. For instance, we noticed that the defence marker genes *PR5* and *PR2* were upregulated, as was *PR1*, the classical marker gene for salicylic acid (SA) defences and SAR (Ryals *et al*., 1996; van Loon *et al*., 2006; **Table S5**). Downregulated processes included primarily photosynthesis, metabolisation of tetrapyrroles, responses to auxin and genes related to decreased oxygen and hypoxia (**Fig. 4b**, **Table S7**). Taken together, the shoot transcriptomes revealed that plants growing on BX_plus_ soil were developmentally more advanced, i.e. towards senescence and with reduced photosynthesis, while having activated SA defences.

**Figure 4.**
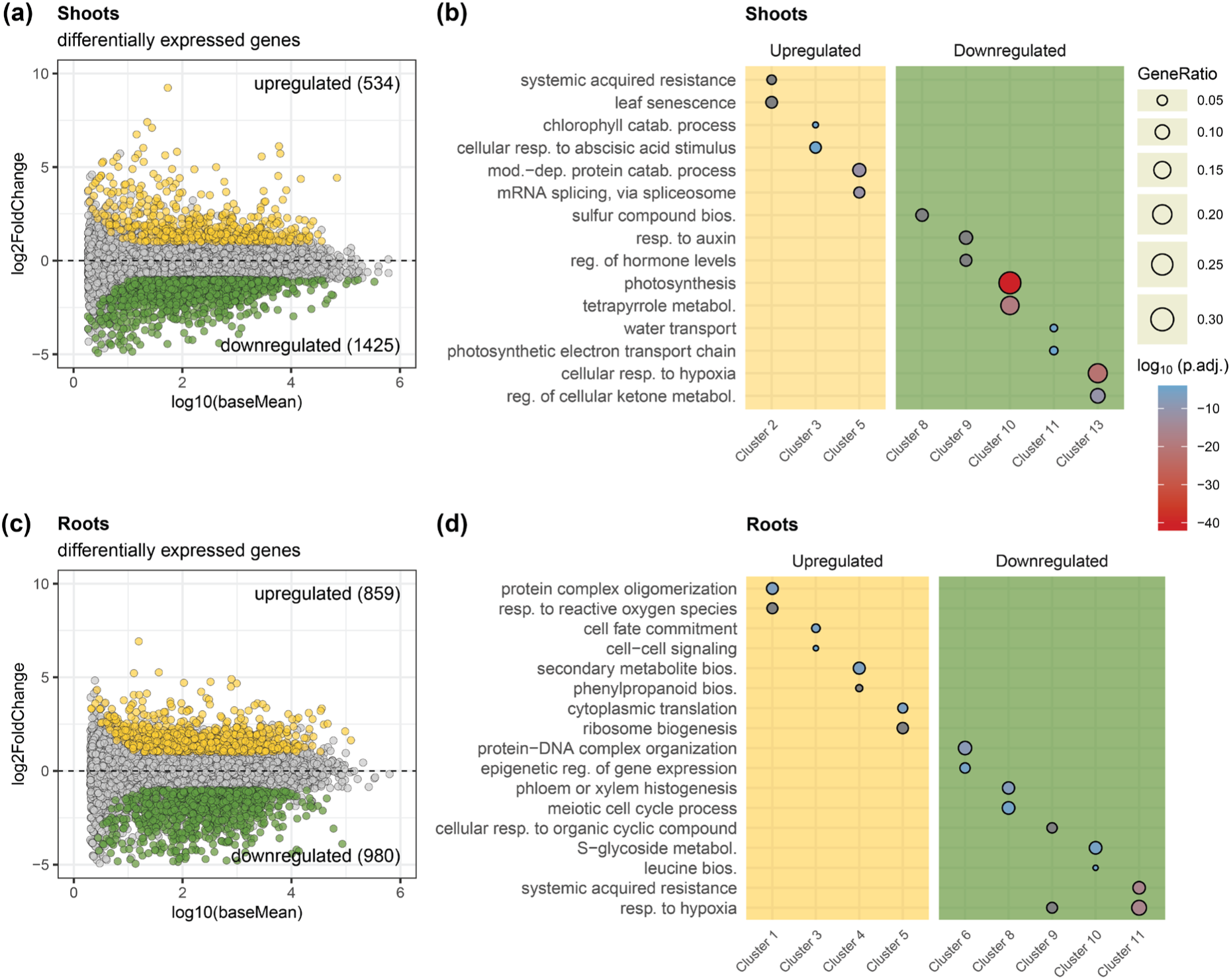
Transcriptome analysis of roots and shoots of Arabidopsis grown on BXplus compared to BXminus soil. **(a)** MA plot of genes expressed in shoots, plotted by their expression level on the x-axis and by the log2FoldChange (log2FC) between BXplus and BXminus soils on the y-axis. Dots in yellow represent genes that were upregulated in shoots grown on BXplus soil compared to BXminus soil, while green dots represent downregulated genes (criteria: *p*adj < 0.05, log2FC > ±1), and grey dots represent genes that were not differentially regulated. The numbers of differentially expressed genes are reported in the MA plot. **(b)** Representation of the top two GO terms in each co-expression cluster for up- and downregulated genes. The top GO terms per cluster were defined by the highest gene ratios, i.e. fraction of differentially expressed genes of a GO term. Colour and size represent the adjusted *p*-values for enrichment of the GO term and its gene ratio, respectively. Some GO terms were shortened due to space (see **Tables S7** and **S8** for full GO terms). **(c)** and **(d)** report the same information for the roots.

In roots, 859 and 980 genes were up- or downregulated, respectively (**Fig. 4c**, **Table S6**). The main upregulated processes included protein complex oligomerization, response to reactive oxygen species, cell fate commitment and signalling, and secondary metabolite biosynthesis (**Fig. 4d**, **Table S8**). The main downregulated processes were related to SAR, responses to low oxygen, S-glycoside metabolism and various developmental processes (**Fig. 4d**, **Table S8**). In summary, the root transcriptomes revealed that plants growing on BX_plus_ soil altered root developmental processes and had downregulated defences. Overall, we concluded that BX-conditioning changed defence processes in both shoots and roots, and also altered growth and development related processes.

### Arabidopsis grown on BX_plus_ soil is better defended against Botrytis and primed for salicylic acid induced defences

The transcriptome analysis revealed a signature of enhanced defences in shoots of plants grown on BX_plus_ soil (**Fig. 4b**). To test actual disease resistance of Arabidopsis, we infected leaves with the necrotrophic fungus *Botrytis cinerea* and quantified the area of the developing lesions. We recorded slightly smaller lesions on plants grown on BX_plus_ compared to BX_minus_ soil in three independent experiments performed on three different soil batches (**Fig. 5a**). Hence, Arabidopsis plants grown on BX_plus_ were more resistant to the pathogenic fungus.

**Figure 5.**
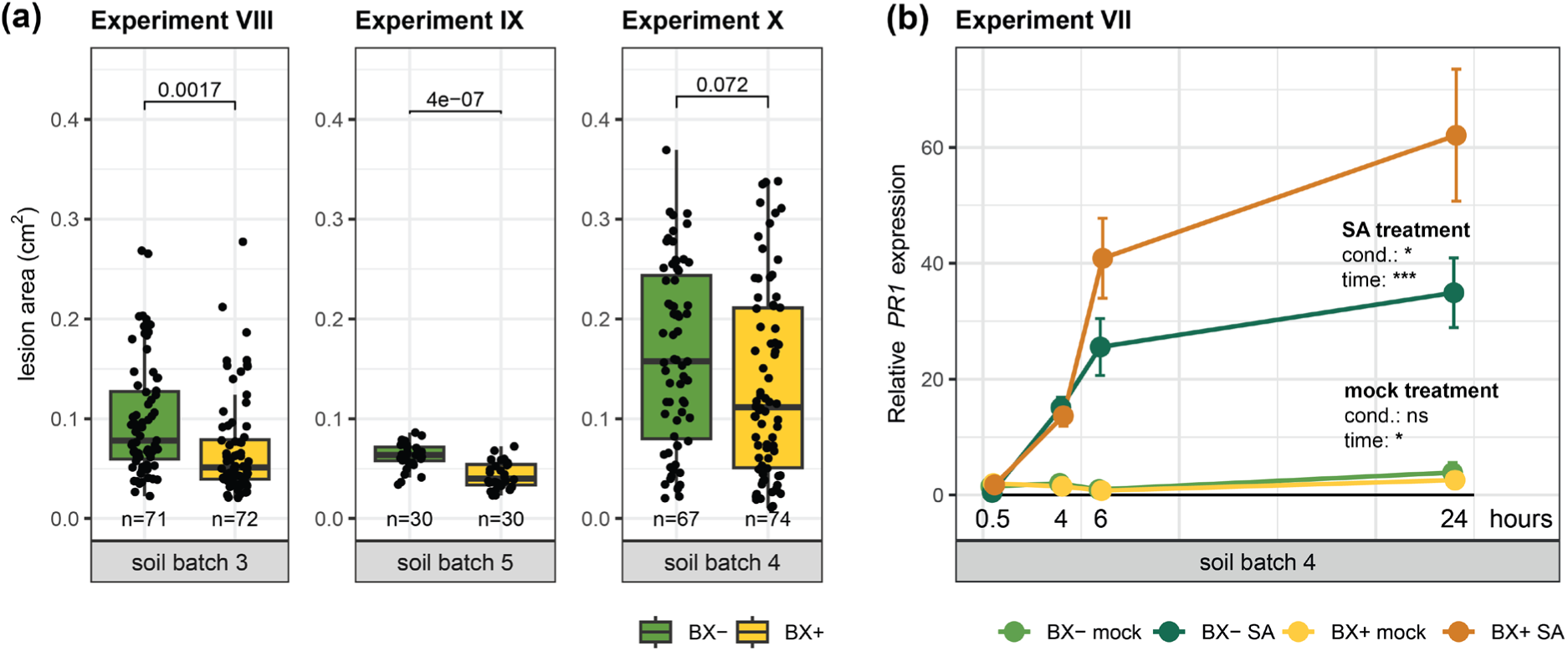
Enhanced resistance of Arabidopsis to Botrytis and primed *PR1* expression on BXplus soil. **(a)** Quantification of *Botrytis cinerea* infections based on the area of necrotic lesions on Arabidopsis leaves. Plant leaves were infected at five weeks for Experiment IX and at six weeks for experiments VIII and X. *P*-values of two-sided student’s t-tests are displayed above each comparison and sample sizes are indicated at the bottom of each boxplot. The soil batch is indicated at the bottom of the graphs. **(b)** Expression of *PR1* relative to the reference gene *PP2A* in mock treated and salicylic acid (SA) treated shoots after 0.5, 4, 6 and 24 hours, in plants grown on BXplus and BXminus soil. The significance levels of ANOVA testing the effects of the conditioning and time after treatment on *PR1* expression are depicted on the right side of the graph, for both treatments separately.

It is formally possible that the enhanced resistance to Botrytis was not triggered by the microbiota but chemically, i.e. by the BX levels in soil (**Fig. S1a**). Thus, we tested whether MBOA additions would affect Arabidopsis’ resistance to the fungus, both in sterile and soil conditions. In a sterile glass jar system (McLaughlin *et al*., 2023), exposure of the plants to 50 µM MBOA in the growth medium decreased not only shoot biomass but also increased lesion area, indicating decreased resistance to Botrytis (**Fig. S5a**). In soil however, exposures of up to 500 µM MBOA did not change resistance to Botrytis, whether added at sowing or only three days before infection (**Fig. S5b**). Therefore, we concluded that MBOA could not trigger Arabidopsis’ resistance to Botrytis, suggesting that residual BXs in soil are unlikely to induce the plant defences directly.

In the shoots, the BX_plus_ soil and/or root microbiota triggered an enhanced expression of genes involved in SAR (**Fig. 4b**). The phenomenon SAR is mediated by the phytohormone SA and establishes after survival of a primary pathogen infection through primed defences (Zeier, 2021). Priming refers to stronger and/or faster defence gene induction upon a second challenge (Martinez-Medina *et al*., 2016). To test whether Arabidopsis’ defences were primed when plants were grown on BX_plus_ compared BX_minus_ soil, we treated the shoots of plants grown on both soils with SA (as secondary challenge) and recorded *PR1* expression levels over time. While *PR1* was not induced in mock-treated plants from both soils, following the SA treatment we found increased *PR1* expression in plants grown on BX_plus_ compared BX_minus_ soil (**Fig. 5b**). Combining these findings allowed to conclude that the BX_plus_ soil and/or root microbiota primed the SA-dependent defences, which is consistent with Arabidopsis’ enhanced resistance to Botrytis when grown on BX_plus_ compared to BX_minus_ soil.

## Discussion

Plant-soil feedbacks (PSFs) describe the performance of a new plant generation to the soil legacy caused by the previous plant generation. We previously reported PSFs where maize growth leads to BX- dependent alterations of the soil microbiota, which then drives the feedbacks of next generation maize or wheat plants (Hu *et al*., 2018; Cadot *et al*., 2021). Mechanistically, how plants respond to altered soil microbiotas and in turn modulate their own performance is not well understood so far. In this study, we established a model system with Arabidopsis to investigate its feedback responses to differential microbiotas in soil, which were established by differential BX exudation of maize. With **Figure 6** we summarize our work and discuss below the implications of our findings in the broader context of how plants mediate microbiome feedbacks.

**Figure 6.**
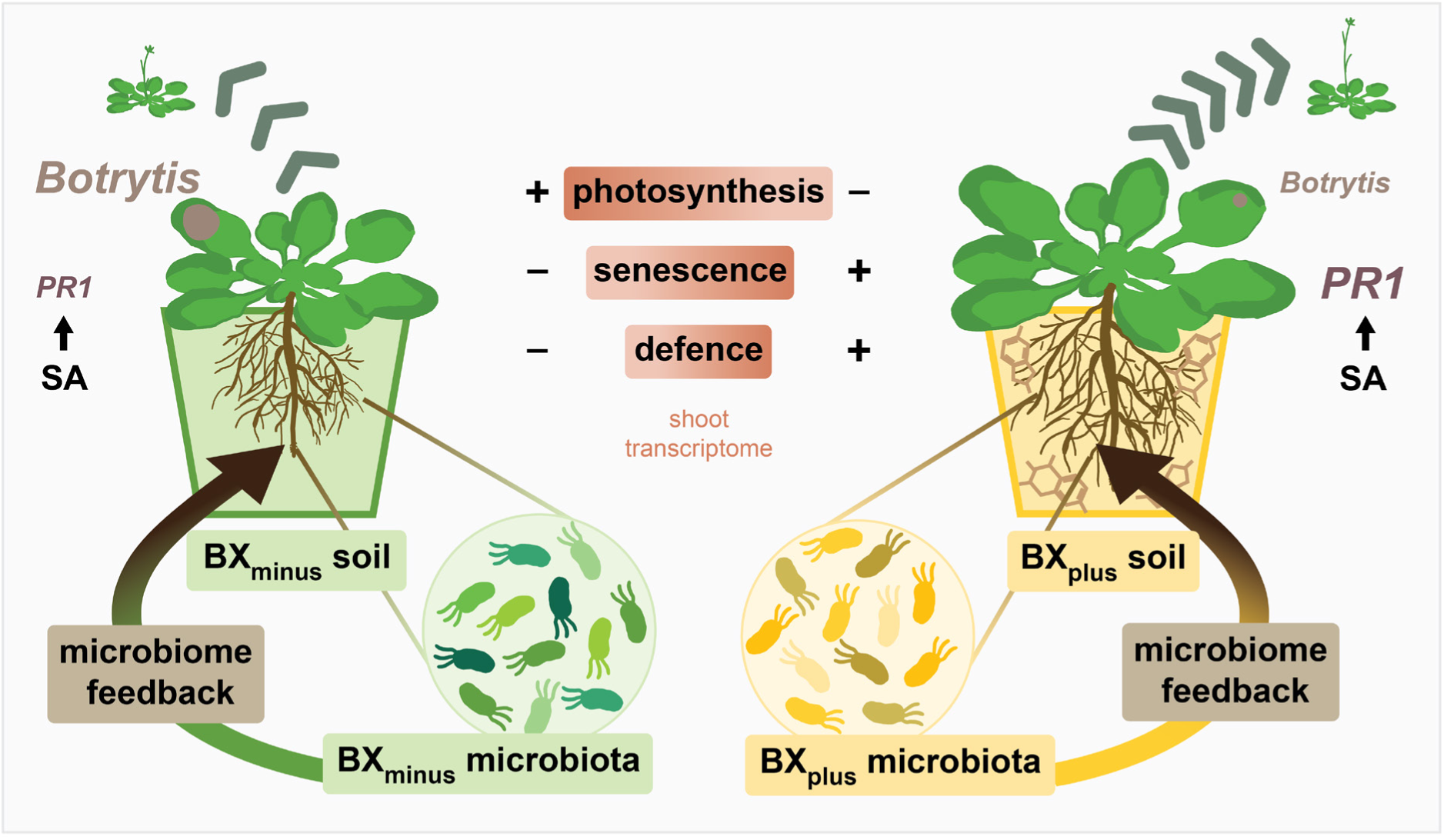
BX-dependent microbiome feedbacks in Arabidopsis. Graphical summary of Arabidopsis responses when grown on BXplus soil compared to the control soil BXminus. BXminus and BXplus soils are physico-chemically the same, but differ in previous conditioning by either wild-type benzoxazinoid (BX) exuding maize or non BX-exuding mutant lines, respectively. The exudation of BXs to soil (indicated with chemical structures in the BXplus pot) affects the soil and root microbiota compositions. Arabidopsis grown on BXplus soil increased root and shoot growth, accelerated development (indicated with the green arrows and icons of bolted plants), was primed for *PR1* and was better defended against *Botrytis cinerea* (lesion sizes are indicated) compared to Arabidopsis grown on BXminus soil. These phenotypic feedbacks were supported by shoot transcriptome analyses (brown boxes with colour gradient), which revealed that plants on BXplus soil were developmentally more advanced, with reduced photoynthesis and increased senesence processes, and that they had enhanced defences. The positive growth on BXplus soil was lost after soil sterilisation, revealing that the differential microbiotas shape the feedbacks, which we therefore, refer to them as microbiome feedbacks (depicted by the arrows leading from the microbiota back to the plant).

In summary, we found that Arabidopsis responded with increased shoot and root growth, faster development and also SA-dependent priming and enhanced resistance to shoot infections with the pathogen *Botrytis cinerea* when grown on BX_plus_ soil (**Fig. 6**). These BX-feedback phenotypes were consistent with the transcriptome analysis revealing also that plants were developmentally more advanced (earlier senescence, reduced photosynthesis) while having enhanced defences in shoots when grown on BX_plus_ soil. We found differential root microbiotas when Arabidopsis was grown on the two soils and also that the positive growth feedback was lost after soil sterilisation. Both observations are consistent with earlier work on maize (Hu *et al*., 2018), corroborating the conclusion that the feedbacks are dependent on BXs but driven by soil microbes. The discovery of the plant immune receptor Mediator of Microbiome Feedback 1 (MMF1) in Arabidopsis feedbacks to BX_plus_ soil (Janse Van Rensburg *et al*., 2025) implies recognition of microbial signals from soil, providing additional evidence that the feedbacks are caused by the soil microbiota. Hence, BX-feedbacks are microbiome feedbacks (**Fig. 6**). The occurrence of such feedbacks with Arabidopsis with its many available research resources make this a good model system to study the molecular mechanisms of microbiome feedbacks.

### Chemical or microbiome feedbacks?

The finding that the better growth of Arabidopsis on native BX_plus_ soil was reversed on sterilised soils (**Fig. 2a**) requires a detailed reflection. While the soil microbiota was eliminated by sterilisation of the soil, the BX contents stayed intact (**Fig. S1a**), representing the main chemical difference between BX_minus_ and BX_plus_ soil (Hu *et al*., 2018; Gfeller *et al*., 2023b). Intuitively, given the allelopathic nature of BXs, this negative feedback on sterilised soil might be due to the residual BXs in soil. Here, we found that Arabidopsis root and shoot growth was only inhibited by MBOA at concentrations in the µM to mM range both in sterile systems or on natural soil (**Fig. 2bc**, **Fig. S2**). Similarly, also AMPO was shown to inhibit Arabidopsis growth in the high µM range (Venturelli *et al*., 2015). In contrast, the concentrations measured in soil were in the nM range (**Fig. S1a**) and thus three or more magnitudes lower. Therefore, it seems unlikely that eventual growth inhibition by soil BXs would explain the reduced growth on sterilised BX_plus_ compared to growth on sterilised BX_minus_ soil.

Alternatively, the negative growth feedbacks on sterilised soils might also be mediated by the newly established microbiota. In support of this hypothesis, the bacterial communities also differed by BX-conditioning on roots from sterilised soils (**Fig. 3c**) and possibly they might explain the differences in shoot growth. These different communities also imply that the residual BXs in sterilised BX_plus_ soil were strong enough determinants to also differentiate these root communities that had freshly assembled via re-colonization of the soil during Arabidopsis growth. Therefore, we concluded that the chemical component of BX-feedbacks with Arabidopsis plays a more indirect role, and that BX-feedbacks are primarily driven through interactions with the soil microbiota that then affect plant growth.

### Developmental advance of plants grown on BX_plus_ soil

The goal of the transcriptome analysis was to investigate how Arabidopsis achieves the better growth on BX_plus_ soil. While no typical growth-related processes were upregulated, we found indications that plant development could be affected: photosynthesis related genes were downregulated and senescence processes upregulated in plants grown on BX_plus_ soil (**Fig. 4b**). Assessing developmental traits, we found indeed that Arabidopsis had more leaves at late growth stage (**Fig. S1e**), showed earlier bolting (**Fig. 1c**) and also produced more seeds (**Fig. 1d**) when grown on BX_plus_ soils. Collectively, this allowed to conclude that plants grown on BX_plus_ soil were developmentally more advanced. A possible explanation for the developmental advance could be differences in germination, as previous work with wheat showed better germination on BX_plus_ compared to BX_minus_ soil (Gfeller *et al*., 2023b). Future experiments are needed to investigate germination effects and to disentangle them from developmental processes involved in BX-feedbacks.

The earlier transition to flowering (**Fig. 1c**) is particularly interesting coupled with the finding of a strongly enriched and abundant Massilia bacterium (bASV39) on roots of plants grown in BX_plus_ soil (**Fig. 3f**). Massilia bacteria are recurrent members of plant root microbiomes including Arabidopsis and they are often associated with beneficial traits (Beilsmith *et al*., 2021; Yu *et al*., 2021; Li *et al*., 2025). Massilia was also recently reported to drive growth promotion of maize and that this beneficial trait depended on flowering time (Wang *et al*., 2024). Hence, Oxalobacteraceae and particularly Massilia clearly warrant future functional investigations with isolates of this genus for their involvement in BX-feedbacks on Arabidopsis.

### BX_plus_ grown plants have enhanced defences

Previously, we have reported better resistance of maize to herbivores when grown on BX_plus_ compared to BX_minus_ soil (Hu *et al*., 2018; Cadot *et al*., 2021). In this study, we found increased resistance of Arabidopsis to the necrotrophic pathogen *Botrytis cinerea* when grown on BX_plus_ soil (**Fig. 5**). In maize, the increased resistance coincided with higher levels of the defence hormones salicylic acid (SA) and jasmonic acid (JA), and most importantly with increased expression of JA-related defence marker genes in the shoots, even when plants were unchallenged (Hu *et al*., 2018). Thus, increased JA-related defence markers on BX_plus_ soil might have facilitated resistance to caterpillars, as defence against herbivores and necrotrophic pathogens mainly requires the JA-related defence pathway (Shigenaga & Argueso, 2016). In Arabidopsis however, we found SA-dependent defences to be upregulated in the shoots of Arabidopsis grown on BX_plus_ soil, as evidenced with the increased expression of typical SA-dependent marker genes including *PR1*, *PR2* and *PR5* (**Fig. 4b**, **Table S5**). Also, treatment with SA lead to higher *PR1* levels in shoots of BX_plus_ grown plants (**Fig. 5b**). As *PR* genes are also upregulated during Botrytis infection (Govrin & Levine, 2002), higher *PR* expression levels on BX_plus_ soil could mediate the increased resistance of Arabidopsis to Botrytis.

Taken together, maize and Arabidopsis differ in the type of hormonal signalling, but both plant species share the commonality of enhanced activation of defence genes when mediating the microbiome feedbacks. This shared defence gene activation appears a possible molecular target for translation. Based on comparative analyses of growth-regulatory mechanisms, Inzé and Nelissen proposed that the particular focus on gene space conservation will enhance the translatability of genetic networks from model to crop species (Inzé & Nelissen, 2022). In the context of the microbiome feedbacks on BX_plus_ soil, thus, future work targeting for example conserved transcription factor families that could mediate defence gene activation both in Arabidopsis and maize seems a logic step forward.

### The BX_plus_ microbiota primes for enhanced defences?

The increased *PR1* expression after SA treatment in plants grown on BX_plus_ soil implies that these plants were primed. Priming is a state of alert where plants can mount a faster and stronger defence response than un-primed plants (Martinez-Medina *et al*., 2016; Karasov *et al*., 2017; Harris *et al*., 2023). To establish defence priming in plants, a preceding priming stimulus is needed, such as a defended pathogen attack, which is described as systemic acquired resistance (SAR; Ross, 1961; Zeier, 2021), or root colonization by beneficial microbes, which is known as induced systemic resistance (ISR; Pieterse *et al*., 1996, 2014). ISR and SAR are typically studied with single beneficial or pathogenic strains, but there is emerging evidence that also whole soil microbiotas can elicit enhanced resistance for example against herbivores (Badri *et al*., 2013; Hubbard *et al*., 2019; Pineda *et al*., 2020). Hence, it appears plausible that priming via ISR or SAR, or even via both processes, can occur in complex microbiotas as it is the case with the differential BX_plus_ and BX_minus_ microbiotas in our study. As we have shown that MBOA itself does not increase Arabidopsis resistance to Botrytis (**Fig. S5**), we concluded that the BX_plus_ microbiota is responsible for priming of *PR1* and for the increased resistance to Botrytis.

Alternatively, the observed enhanced resistance might be due to direct microbe-microbe interactions on the leaves, as it was recently shown for the soil legacies established by downy mildew infected Arabidopsis, where disease suppressive bacteria from soil enriched on Arabidopsis leaves and inhibited downy mildew sporulation (Goossens *et al*., 2023). Future work should thus clarify how resistance to Botrytis is established in Arabidopsis grown on BX_plus_ soil. However, as priming usually exerts no or only little costs compared to constitutive defences, it could serve as the explanatory mechanism for the lack of a growth-defence trade-off, which was observed with the improved growth and the enhanced resistance of Arabidopsis on BX_plus_ soil.

### Relevance of Arabidopsis BX-feedbacks

In this study, have found enhanced growth of Arabidopsis on BX_plus_ compared to BX_minus_ soil (**Fig. 1**), which is a positive growth feedback. In contrast, neutral to negative feedbacks on maize growth and positive to negative feedbacks on wheat growth were reported, depending on the genotype and field soil (Hu *et al*., 2018; Cadot *et al*., 2021; Gfeller *et al*., 2023b,a). These observations suggest that the varying feedback direction – positive or negative – depends both on the soil environment and on the genotype of the response plant. For the former, we have demonstrated that BX-feedbacks can vary in strength and direction along a soil chemical gradient that exists within a tested field in Changins (Gfeller *et al*., 2023a). Future work is needed to extend on the second part of the above conclusion - whether and how the genotype of the response plant influences the direction of BX-feedbacks. Arabidopsis with its many accessions might be useful in uncovering genetic components of such microbiome feedbacks.

Furthermore, our findings with Arabidopsis highlight that also dicots and plants that - unlike maize or wheat - do not produce BXs themselves, respond to BX-conditioned soil microbiomes. Notably, we report increased growth and resistance to a necrotrophic fungus, while for maize and wheat, often both positive and negative effects were reported simultaneously, such as increased resistance to insects but decreased growth for the maize variety B73 (Hu *et al*., 2018). This is not surprising, as ecological studies revealed that con-specific PSFs are often negative (Jing *et al*., 2022), especially for con-specific but also inter-specific feedbacks in the Poaceae family (Hannula *et al*., 2021). Following the framework of bridging knowledge from PSFs in natural and agricultural systems (Mariotte *et al*., 2018), the potential of BX-feedbacks to elicit positive but not negative feedbacks in more distant crops such as rapeseed (*Brassica napus*) should be further explored. Brassicaceae crops are often used in rotation schemes because of their beneficial traits of glucosinolates functioning in biofumigation (Gimsing & Kirkegaard, 2009). Field work testing a rotation where rapeseed follows maize appears the logical next step to investigate the translatability of the microbial feedbacks studied with the Arabidopsis (also a Brassicaceae) model system established in this study.

## Conclusion

The potential to utilize the beneficial functions of the soil or root microbiome as a more sustainable way to improve plant growth and health in agriculture is well recognized (French *et al*., 2021; Santos & Olivares, 2021). However, the mechanisms by which plants perceive and respond to specific soil microbiomes are currently not well understood (Janse van Rensburg *et al*., 2024). Our study provides a first step into this direction, as we show that Arabidopsis can respond with both improved growth and enhanced resistance on a specific soil microbiota. The model system with BX-conditioned soil microbiomes and their feedbacks on Arabidopsis will allow us to further study the mechanisms by which Arabidopsis can perceive and mediate microbiome feedbacks, and especially to identify the plant genes that are important for this process. Possibly, such knowledge can help to improve breeding programs so that crops can take full advantage of their microbiome.

## Acknowledgements

We sincerely thank Nicolas Widmer and Dr. Luca Bragazza (Agroscope Changins, Switzerland) for field access and assistance in soil sampling, Prof. Tobias Züst (University of Zurich, Switzerland) and Mirco Hecht (Federal Statistical Office, Switzerland) for the BX analysis, and Dr. Phillipe Demougin and Dr. Geoffrey Fucile (both University of Basel, Switzerland) for preparation and sequencing of the RNAseq library. Furthermore, we thank Corinne Suter (University of Bern, Switzerland) for implementing the ‘in vitro Experiment I’, Markus Funk (University of Basel, Switzerland) for performing the RNA extractions, Dr. Selma Cadot (Agroscope Reckenholz, Switzerland) for collecting field soil, Jan Wälchli (University of Basel, Switzerland) for assembling the microbiota reads and Florian Enz (University of Bern, Switzerland) for maintenance of maize plants. Lastly, we are also grateful to Prof. Christelle Robert and Prof. Matthias Erb (both University of Bern, Switzerland) for providing the maize seeds, and to Prof. Wim Van den Ende (KU Leuven, Belgium) for providing the Botrytis strain. This work was mainly supported by fundings of the University of Basel and partly by the University of Bern (Interfaculty Research Collaboration “One Health” to K.S.) and the Swiss National Science Foundation (No. 189249 to K.S.).

## Author contributions

KSt, LT and KS planned and designed the research. KSt, HJvR, LT and VD performed experiments. KSt and LS analysed the data. KSt and KS wrote the manuscript.

## Data availability

The source data on plant biomass and soil physicochemical analyses is made publicly available on https://github.com/PMI-Basel/Stengele_et_al_At_BX-feedbacks. The raw sequencing data are stored at the European Nucleotide Archive (http://www.ebi.ac.uk/ena). The RNA Seq sequencing data is stored under project number PRJEB80860, and the microbiota sequencing data is available under the project number PRJEB59165 (The 16S rRNA and ITS gene libraries of this study were sequenced in the same MiSeq run together with a library of an already published project (Gfeller *et al*., 2023a)). All bioinformatic code including information on barcodes, primers and sample assignments is provided on GitHub. On GitHub we have also deposited all code used for statistical analysis and graphing of all figures.

## Supplementary Methods

### Feedback experiments on soil

We have performed several experiments (Exp. I to Exp. XII) with slightly different setups, which are detailed in **Table S2**. While the general procedure, which applies to all experiment, is described in the main text, here we explain here additional details of particular experiments:

*Experiment III:* Fractions of the soil/sand mixtures for BXplus and BXminus soil were additionally sterilised by X-radiation (36 – 40 kGy, Synergy Health AG, Däniken, Switzerland) before filling to pots. Also, seeds for Experiment III were sterilised (Protocol, see in vitro experiment 2) before sowing and pots were watered with autoclaved tap water for the whole course of the experiment.

*Experiment VII:* The number of leaves was counted from images of the rosettes taken at different time points. *Experiment XI:* This experiment, testing the application of MBOA to soil, was conducted with unconditioned soil collected from the field in Changins and was performed testing ‘long-term’ and ‘short-term’ exposure. One group of control pots remained untreated and only received water for the duration of the experiment. Half of the treated (DMSO and different concentrations of MBOA (dissolved in DMSO)) pots were used for ‘long- term’ exposure to MBOA and they were treated before sowing. The other half of the treated pots were used to test ‘short-term’ exposure and they were treated after six weeks of plant growth (and three days before infection with Botrytis; see *Infections with Botrytis cinerea* in the main Materials and Methods). The treatments con- sisted of applications of 12 mL of tap water containing 0, 100 or 500 µM MBOA in 0.06% DMSO. The rosette area data reported in this manuscript for Experiment XI were of ‘long-term’ exposed plants, while the Botrytis infection was performed on both ‘long-term’ and ‘short-term’ exposed plants.

*Experiment XII:* In this experiment, plants had been grown in a climate chamber (Sanyo, Moriguchi, Japan) equipped with fluorescent bulbs under long-day conditions (16 h day at 21°C and light intensities between 100 and 200 µmol m^-2^ s^-1^, and 8 h night at 18°C). To score feedbacks on flowering, we recorded at three time points the following three developmental stages: plants (i) did not bolt yet, (ii) showed onset of bolting and (iii) had bolted. For the plants that had bolted, we further measured the heights of the developing flower stalks using a ruler. Plants that had not bolted at the third timepoint were discarded, and the rest of the plants were grown to seed, and we collected the seeds for each plant separately (n=12 on each soil). Seed yields were scored by measuring the total seed weight on an analytical balance and by spreading the seeds on a grid paper followed by manually counting them using a mechanical tally counter. Finally, we calculated the average weight of 100 seeds based on the total seed number and total seed weights.

### In vitro experiments

*Experiment 1:* The seeds were sterilised for four hours with chlorine gas (Lindsey III *et al*., 2017), and then directly sown to ½ Murashige and Skoog (MS; Duchefa, Haarlem, the Netherlands) plates containing 1.5% plant agar (Duchefa, Haarlem, the Netherlands) supplemented with MBOA (Sigma-Aldrich, St. Louis, USA) at different concentrations (0, 100 or 500 µM) or no MBOA as control. MBOA was dissolved in DMSO (Sigma-Aldrich), which was kept constant at 500 µL per L ½ MS agar in each treatment including the control. 12 to 16 seeds were sown per plate and stratified for three days at 4°C. Seed germination was recorded for the first four days after germination, and primary root length was quantified ten days after sowing.

*Experiment 2*: Seeds were surface sterilised with 70% Ethanol + 0.1% Triton-X100 for 1 minute, hypochlorite solution (<5%, Potz, Migros, Zurich, Switzerland) + 0.1% Triton-X100 for 12 minutes, and washed three times with sterile milliQ water. Seeds were sown onto ½ MS agar plates containing 1% sucrose, stratified for three days at 4°C and germinated for eight days. The seedlings were then transferred to ½ MS plates containing 825.75 µL DMSO per L ½ MS agar as control or plates containing the same amount of DMSO with different MBOA concentrations (0, 50, 100 or 500 µM). Five seedlings were transferred onto each plate and grown for additional 13 days, before the total root and shoot material per plate was harvested to determine the fresh weight.

*Experiment 3:* This experiment was conducted in a semi-hydroponic growth system (McLaughlin *et al*., 2023). Seeds had been sterilised as described for in vitro experiment 2, and grown on ½ MS for seven days. Seedlings were then transplanted to fresh ½ MS plates and grown for another 14 days. Two seedlings were then trans- planted to the jar system described in McLaughlin *et al*. (2023), but with a modified volume of 32.5 mL ½ MS per jar. The ½ MS added to the jars contained final concentrations of either 0 or 50 µM MBOA in 0.1% DMSO. Every nine days of growth in the jars, the growth medium was replaced with fresh 32.5 mL of ½ MS including freshly prepared MBOA or control treatments. After a total of 28 days in the jars, one plant was harvested from the jar for sample material collection. One day later, the remaining plant was used for *Botrytis* infection (see *Infections with Botrytis cinerea* in the main Materials and Methods). After infection, the lid was taped to ensure high humidity for infection. Jars were then placed into growth chambers equipped with LED light under low light conditions (∼ 50 µmol m^-2^ s^-1^) and the lesion area was then quantified three days after infection.

### Microbiota profiling

*Sampling:* The soil samples were taken from the middle of unplanted pots that had been included in experiment III, and a subsample of approximately 250 mg was used for DNA extraction. For roots, we harvested them from the pot and sampled a 5 cm long root fragment starting from 1 cm below the rosette base, which included both primary and lateral roots. These root segments were washed in 10 mM MgCl_2_, blot dried with tissue paper, snap frozen in liquid nitrogen, and stored at −80°C. Root samples were then lyophilized (FreeZone Plus; Labconco Corporation, Kansas City, USA) for three days and ground for 5 minutes with a ball mill (Mixer Mill MM 400; Retsch GmbH, Haan, Germany).

*DNA extraction:* From all samples, we extracted the DNA with the NucleoSpin Soil DNA extraction kit (Ma- cherey-Nagel, Düren, Germany), and quantified the DNA concentrations with the AccuClear® Ultra High Sensitivity dsDNA Quantitation Kit (Biotium, Fremont, USA). We normalized the concentration to 0.2 ng/µL for soil samples and 2 ng/µL for root samples, or used undiluted samples if DNA concentrations were lower. *PCR and library preparation:* We performed two-step PCR amplification and barcoding of the bacterial 16S rRNA gene region and the fungal ITS spacer region as was previously described in Gfeller et al., (2023b). We used the same cycling profiles, but with 3 minutes of initial denaturation. The reaction of the first 16S rRNA gene PCR was composed of 1x 5Prime HotMasterMix (Quantabio, Beverly, USA), 0.3% BSA, 300 nM of each primer and 10 ng root DNA or 1 ng soil DNA. The second PCR contained 6 µL of purified PCR product (using SPRIselect beads; Beckman Coulter Life Sciences, Indianapolis, USA) and individual barcoded primer pairs for each sample. The ITS PCR reactions contained 200 nM of each primer but were otherwise the same as the 16S rRNA gene reactions. The products from the second PCR were then purified with SPRIselect beads, and pooled to 25 ng DNA per sample for the 16S samples, and to 2.5 ng DNA or as much as was available per sample for the ITS samples. Both pools were bead purified again and pooled for the final library.

*Sequencing and bioinformatic analysis:* The library was sequenced with the MiSeq reagent kit v3 at the Next Generation Sequencing Platform (University of Bern) using the 2x 300 bp pair-end sequencing protocol (Illu- mina Inc., San Diego, USA). The MiSeq 16S rRNA gene and ITS reads were quality checked with FastQC v0.11.8 (Babraham Institute, Cambridge, United Kingdom) and demultiplexed with cutadapt v2.10 (Martin, 2011). Quality filtering, read merging and amplicon sequence variants (ASV) clustering was implemented in R v4.0.0 (R Core Team, 2017) using the package dada2 v1.16.0 (Callahan *et al*., 2016). Bacterial taxonomies were assigned with a naïve Bayesian classifier using a DADA2 formatted training set (silva_nr_v132_train_set.fa.gz) from the SILVA database (Quast *et al*., 2013), and fungal taxonomies were assigned with a training set (sh_general_release_dynamic_02.02.2019.fasta) from the UNITE database (Abarenkov *et al*., 2024).

## Supplementary Figures

**Figure S1.**
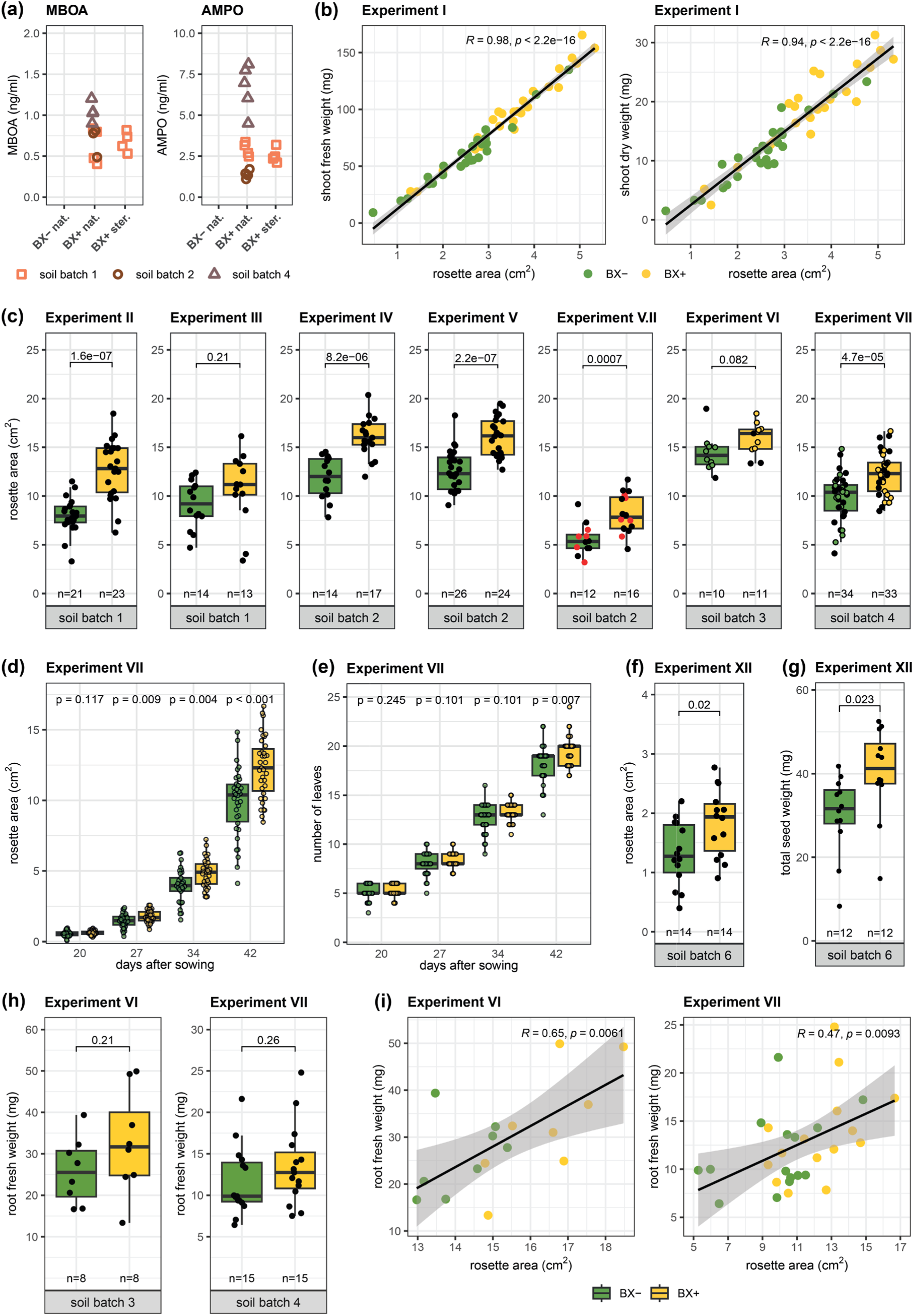
BX concentrations in soils and Arabidopsis growth measurements on different soil batches. **(a)** Concentrations of MBOA and AMPO measured in native as well as sterilised conditioned soils. BX concentrations had been measured in native (‘nat.’) BXplus and BXminus soils for soil batches 1, 2 and 4, and in sterilised (‘ster.’) BXplus soil for soil batch 1; n = 4 for each treatment and soil batch. Samples that were below the detection limit for the given compound are not shown. **(b)** Correlation of Arabidopsis rosette area to shoot biomass (shoot fresh weight and shoot dry weight). The pearson correlation coefficient and respective *p*-value for the correlation are displayed within the graph. **(c)** Arabidopsis rosette area from seven independent experiments. Note that the shoot area data for Experiment III is also shown in Fig. 2a. For Experiment V.II, the red dots indicate the plant replicates where the shoot and root transcriptome had been sequenced for the RNASeq analysis. For Experiments VI and VII, colored dots (yellow and green) represent plants where the root biomass was quantified, whereas black dots indicate no quantification of roots. The soil batch number, indicated at the bottom of each individual plot, denotes the independent conditioning events of the soil by growing maize. **(d)** Rosette area of plants over time. Note that the 42 day timepoint is also shown in the last panel in (c). Colored dots were chosen for better visibility only. **(e)** Visible number of leaves quantified from the rosette photographs over time. For (d) and (e), the fdr adjusted *p*-values of two-sided student’s t-tests are reported on top of each timepoint. **(f)** Arabidopsis rosette area of Experiment XII from 23 days after sowing. Note that plants from Experiment XII were grown under long-day conditions. **(g)** Total weight of all seeds collected per plant in Experiment XII. The soil batch number of the soil conditioning event is indicated at the bottom. **(h)** Root fresh weights of Arabidopsis from two experiments and their **(i)** correlations with the rosette area data. The pearson correlation coefficients and respective *p*-values are displayed within the plots. In (c), (f), (g) and (h), boxplots report the *p*-values of two-sided student’s t-tests and the replicate numbers of the different experiments.

**Figure S2.**
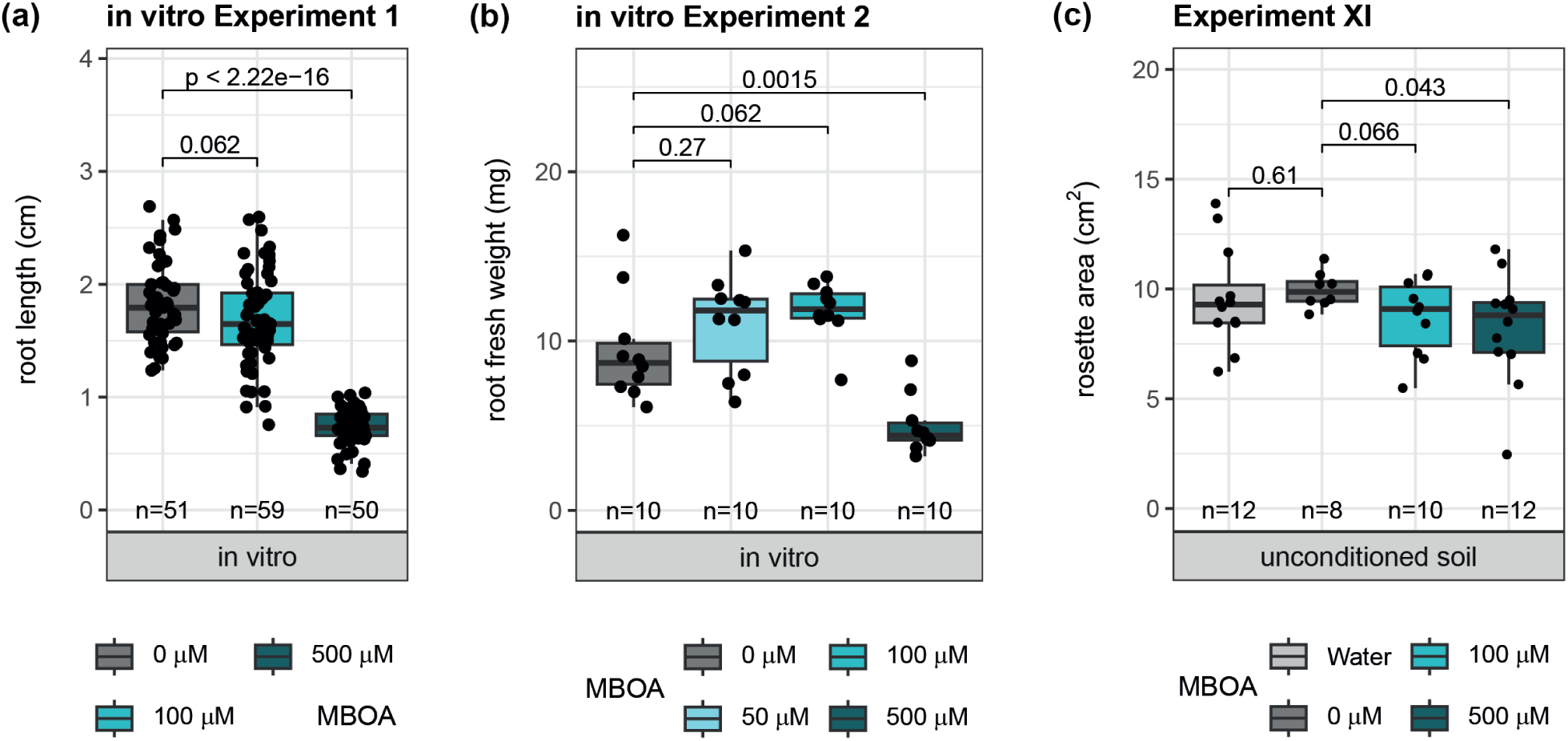
Arabidopsis root growth on MBOA supplemented agar plates and rosette area on soil supplemented with MBOA. **(a)** Root length of Arabidopsis germinated and grown for ten days on ½ MS agar plates supplemented with DMSO (‘0 µm’) or increasing amounts of MBOA dissolved in DMSO. **(b)** Root fresh weight of Arabidopsis grown for 21 days in total. Plants were pre-germinated for eight days on ½ MS agar with sucrose, and then grown for an additional 13 days on ½ MS agar plates supplemented with DMSO or increasing amounts of MBOA dissolved in DMSO. For the root fresh weight data, one data point represents the total root fresh weight from five plants grown on one plate. **(c)** Rosette area of Arabidopsis grown on un-conditioned soil amended with different MBOA concentrations. Before sowing of seeds, pots were either treated with water, DMSO (‘0 µM’) or MBOA dissolved in DMSO. In all graphs, replicate numbers are reported below the boxplots, and the *p*-values of two-sided student’s t-tests are reported for the respective comparisons.

**Figure S3.**
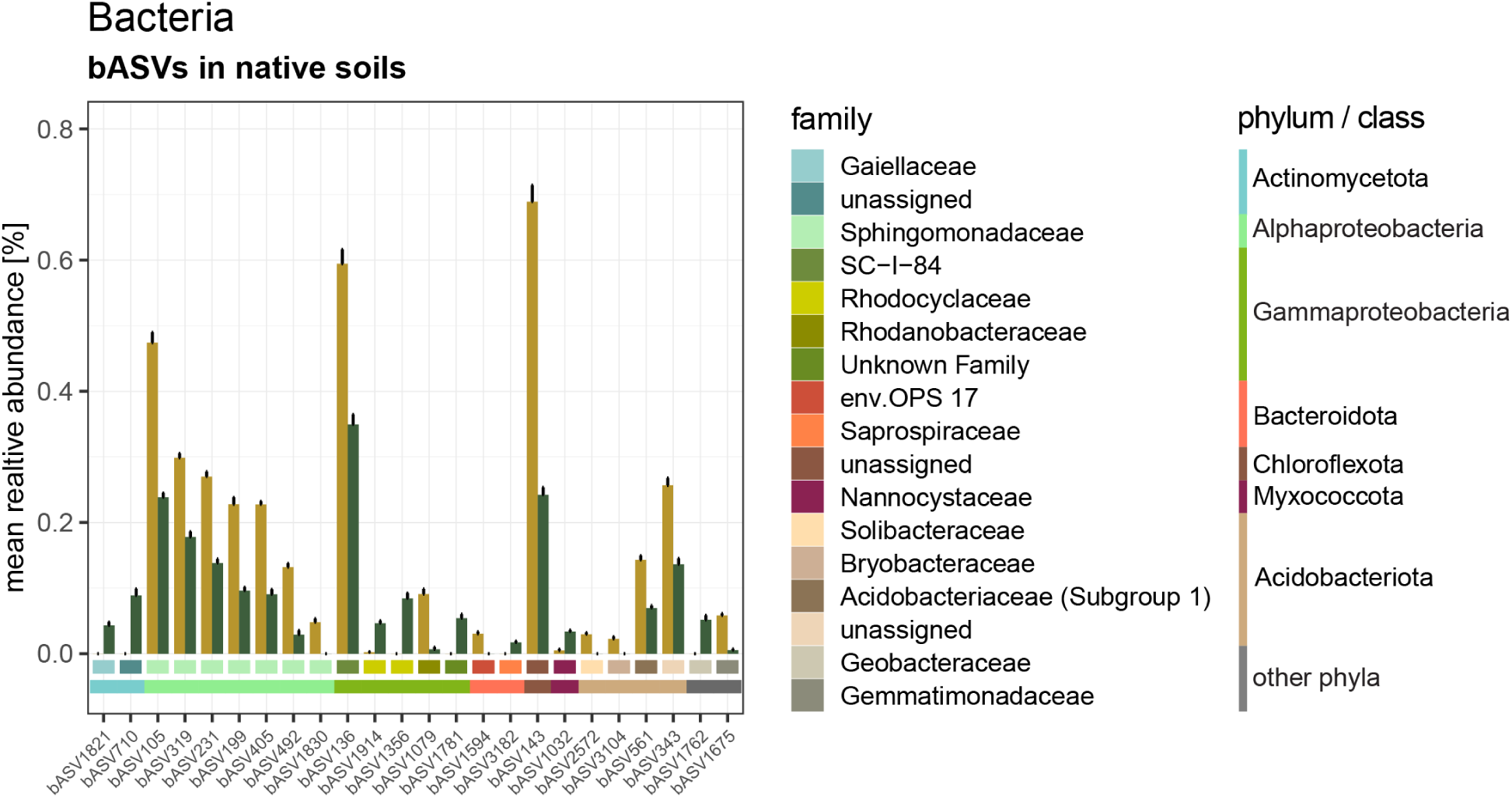
Differential bASVs in native soils. Abundance and taxonomy of differentially abundant bacterial ASVs (bASVs) in native BXplus (brown bars) and BXminus (green bars) soil. Error bars represent the standard error of the mean. The colored horizontal bars below the bargraphs indicate the bacterial family and phylum/class information (for the Pseudomonadota phylum, the class information is shown, ie. Alphaproteobacteria or Gammaproteobacteria).

**Figure S4.**
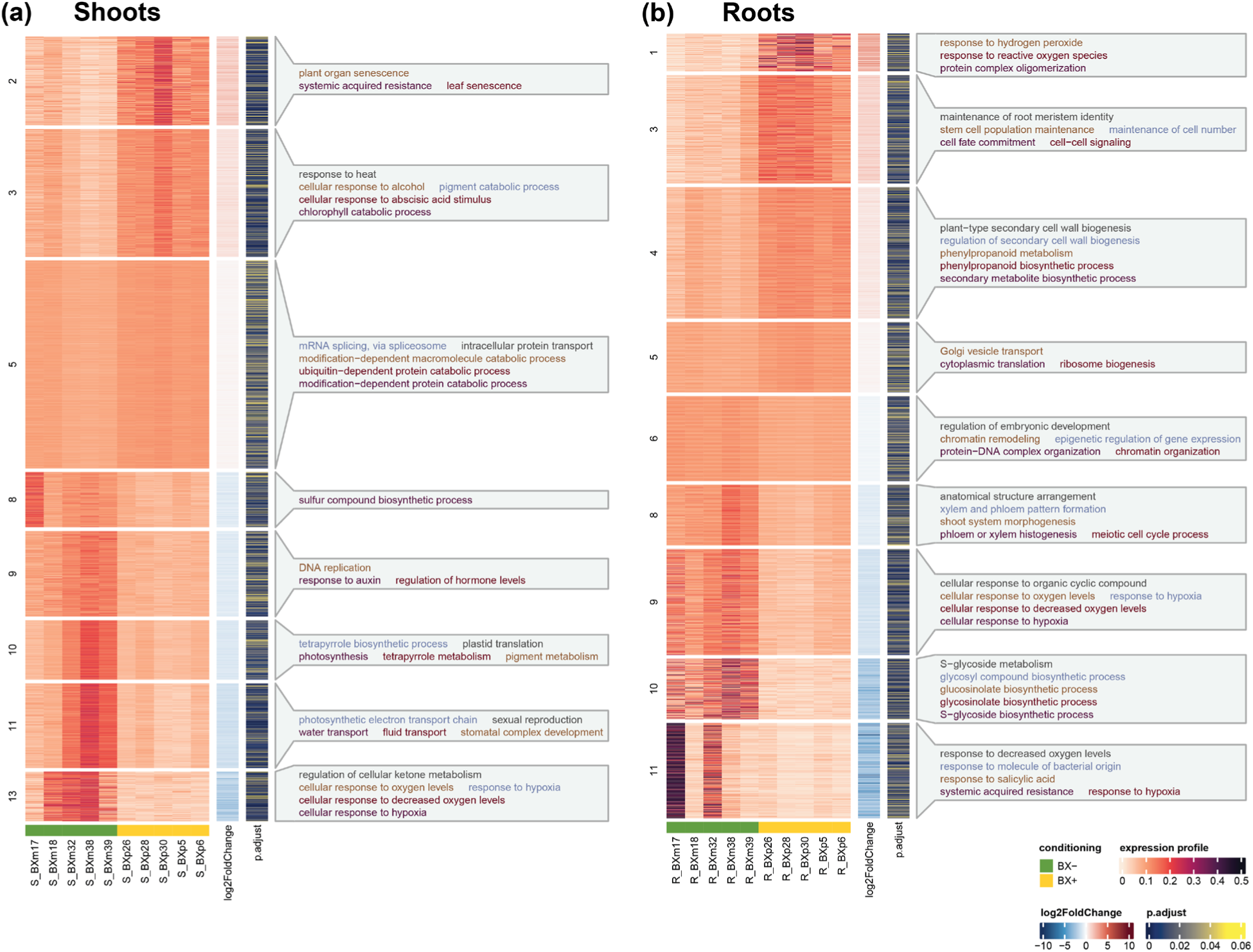
Co-expression analysis of Arabidopsis roots and shoots. Heatmaps of co-expressed gene clusters in **(a)** roots and **(b)** shoots, as identified with coseq. Colors in heatmap correspond to expression profiles i.e., the proportion of normalized counts per gene. Adjacent color bars show the log2FoldChange in expression, as well as the corresponding ajusted *p*-values as inferred with DESeq2. Cluster labels show the top five enriched GO terms (biological processes, ajusted *p*-value < 0.01) in the corresponding clusters. The different GO term text colors are for better readability only. Full tables of all enriched GO terms in each cluster in roots and shoots are provided in **Tables S7** and **S8**.

**Figure S5.**
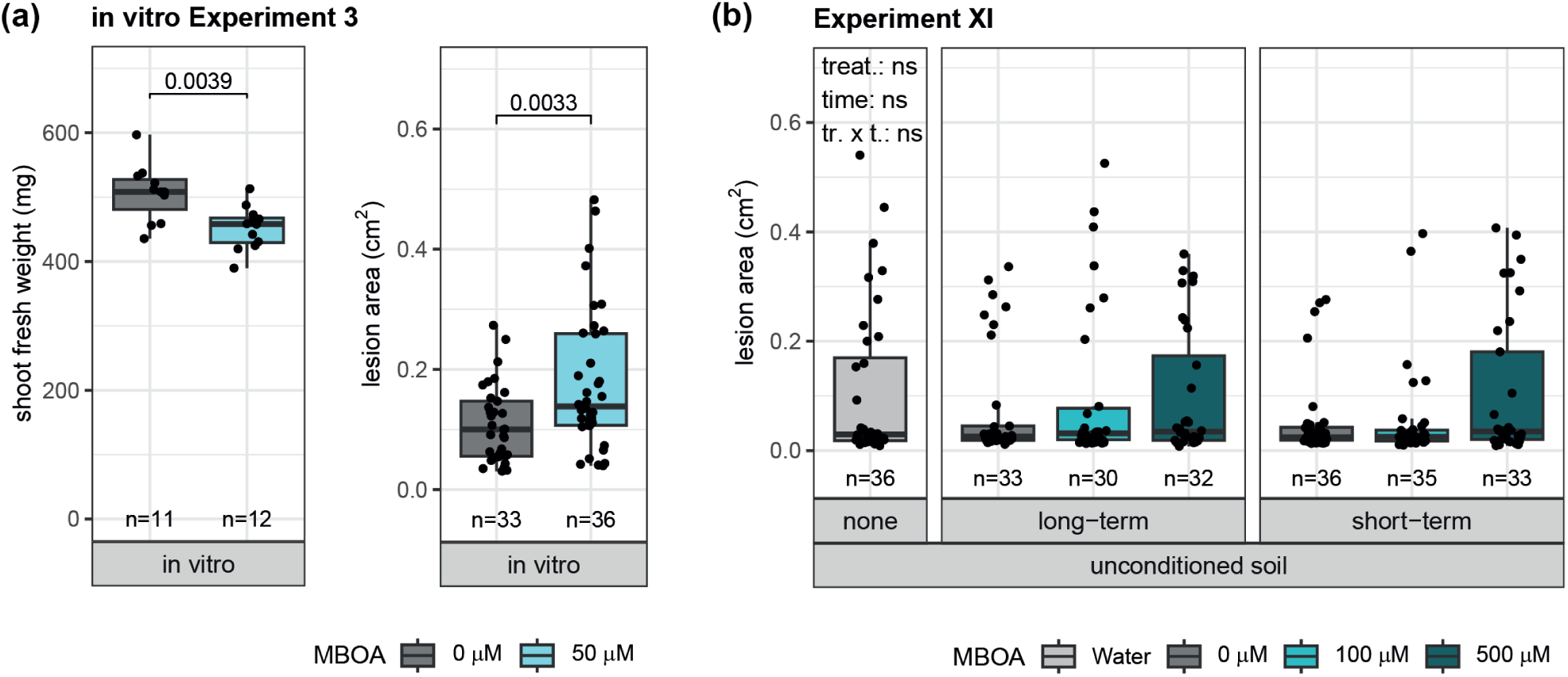
MBOA does not induce enhanced resistance in Arabidopsis against *Botrytis cinerea*. **(a)** Shoot biomass and *Botrytis cinerea* (‘Botrytis*’*) infection of Arabidopsis grown in a semi-hydroponic glass jar system. Plants had been pre-grown for a total of 21 days on sterile ½ MS plates before they were transplanted to glass jars. In the jars, plants were grown in ½ MS nutrient solution treated with DMSO (‘0 µm’) or 50 µm of MBOA dissovled in DMSO. After 36 days of total growth, three leaves per plant were infected with Botrytis, and the lesion area was quantified three days after the infection (graph on the right). The shoot biomass of the remaining shoot (= minus the three leaves that were infected) was also enumerated (graph on the left). The *p*-values of two-sided student’s t-tests are reported in each graph. **(b)** Botrytis infection of Arabidopsis grown on un-conditioned soil amended with different MBOA concentrations. Plants were either untreated for the whole growth period (‘Water’), treated with DMSO (‘0 µm’) or increasing amounts of MBOA dissolved in DMSO before sowing for a ‘long-term’ exposure, or treated with the same concentrations but only three days before the infection for a ‘short term’ exposure. The significance levels of the ANOVA model testing the effect of the chemical treatment (‘treat’) and the timepoint of treatment application (‘time’) on lesion area are depicted in the top left corner of the graph. In all graphs, replicate numbers are reported below the boxplots.

## Supplementary Tables

**Table S1.** Set up of the different conditioning experiments. One ‘soil batch’ denotes one independent conditioning experiment, where we used parts of the collected soils indicated in the ‘soil collection’ column, and grew wild-type B73 and *bx1*(B73) maize plants for twelve weeks to condition the soil. Soils were collected from three adjacent Agroscope research fields in Changins, Nyon, Switzerland (46°24’00.0"N 6°14’22.8"E), and the ‘soil collection’ column denotes when the soil was collected and from which field (parcel). The two fields had the following cropping history: 2017 alfalfa, 2018 maize, 2019 winter wheat [parcel 29] and 2017 spring wheat, 2018 maize, 2019 sunflower [parcels 30 and 31]. For the fertilisation, the pots received the specified amount of the re- spective nutrient solution once per week, starting with a low iron solution for the first four weeks and increasing to a high iron solution for most of the soil batches. rel. hum. = relative humidity, int. = intensity. Fe = iron.

| Soil batch | Soil collec-<br>tion | Pots | Growth<br>environment | Fertilisation | Growth conditions |
| --- | --- | --- | --- | --- | --- |
| soil batch 1 | May 2019<br>(parcel 29) | 2 L (round) | walk-in<br>chamber<sup>1</sup> | 4 w. low Fe<sup>3</sup>,<br>8 w. high Fe<sup>4</sup> | 16 h day at 26°C, 8 h night at 23°C,<br>50% rel. hum., ~550 $\mu\text{mol m}^{-2} \text{s}^{-1}$ light int. |
| soil batch 2 | May 2019<br>(parcel 29) | 2 L (round) | walk-in<br>chamber<sup>1</sup> | 4 w. low Fe ,<br>8 w. high Fe | 16 h day at 26°C, 8 h night at 23°C,<br>50% rel. hum., ~550 $\mu\text{mol m}^{-2} \text{s}^{-1}$ light int. |
| soil batch 3 | Sept. 2022<br>(parcel 29/30) | 3 L (round) | greenhouse | 4 w. low Fe ,<br>8 w. high Fe | 16 h day at min. 23°C, 8 h night at min.<br>19°C, 300 - 500 $\mu\text{mol m}^{-2} \text{s}^{-1}$ light int. |
| soil batch 4 | Sept. 2022<br>(parcel 29/30) | 3 L (round) | Phytotron<sup>2</sup> | 12 w. low Fe | 14 h day at 22°C, 10 h night at 18°C,<br>60% rel. hum., ~550 $\mu\text{mol m}^{-2} \text{s}^{-1}$ light int. |
| soil batch 5 | Aug. 2020<br>(parcel 30/31) | 3 L (round) | greenhouse | 4 w. low Fe,<br>8 w. high Fe | 16 h day at min. 23°C, 8 h night at min.<br>19°C, 300 - 500 $\mu\text{mol m}^{-2} \text{s}^{-1}$ light int. |
| soil batch 6 | Dec. 2021<br>(parcel 30) | 3 L (round) | greenhouse | 4 w. low Fe,<br>8 w. high Fe | 16 h day at min. 23°C, 8 h night at min.<br>19°C, 300 - 500 $\mu\text{mol m}^{-2} \text{s}^{-1}$ light int. |

<sup>1</sup> Kaelte 3000 AG, Landquart, Switzerland
<sup>2</sup> Phytotron Facility, University of Basel, Switzerland
<sup>3</sup> 100 mL of 0.2% Plantaaktiv Typ K (Hauert HBG Duenger AG, Grossaffoltern, Switzerland) and 0.001% Sequestrene Rapid (Maag, Westland Schweiz GmbH, Dielsdorf, Switzerland)
<sup>4</sup> 200 mL of 0.2% Plantaaktiv Typ K, 0.02% Sequestrene Rapid

**Table S2.** Setups of the different Arabidopsis experiments. For each experiment, the soil batch, pot size and growth length are indicated. For details on the soil batch, see Table S1. Soil batches were stored at 4°C and the soil storage column indicates how long the soils were stored before set up of the experiments (in months), while the growth column indicates how long the Arabidopsis plants were grown. The fertilisa- tion column indicates how many times the plants were fertilised during the growth phase for each experiment. As meas- urements, we quantified shoot growth at the end of the experiment, and for certain experiments we additionally recorded flowering, the root biomass, collected roots for a microbiota analysis, collected root and shoot material for a transcriptome analysis, assessed priming of *PR1* or infected the leaves with the fungus *Botrytis cinerea*. Uncond. = unconditioned.

| Experiment | Soil batch | Pots | Soil storage | Fertili- sation<sup>1</sup> | Growth (weeks) | Measurements |
| --- | --- | --- | --- | --- | --- | --- |
| Exp. I | soil batch 1 | round<sup>2</sup> | 4 | - | 7 | Shoot growth |
| Exp. II | soil batch 1 | square<sup>3</sup> | 5 | 3x | 7 | Shoot growth |
| Exp. III | soil batch 1 | square<sup>4</sup> | 8 | 1x | 6 | Shoot growth, Root microbiota |
| Exp. IV | soil batch 2 | square<sup>3</sup> | 6 | 1x | 6.5 | Shoot growth |
| Exp. V | soil batch 2 | square<sup>4</sup> | 8 | 1x | 8 | Shoot growth |
| Exp. V.II | soil batch 2 | square<sup>4</sup> | 8 | 1x | 7 | Shoot growth, Transcriptome |
| Exp. VI | soil batch 3 | square<sup>4</sup> | 0.75 | - | 7 | Shoot growth, Root biomass |
| Exp. VII | soil batch 4 | square<sup>4</sup> | 2.5 | 2x | 7 | Shoot growth, Root biomass, *PR1* priming |
| Exp. VIII | soil batch 3 | square<sup>4</sup> | 1 | 2x | 6 | Shoot growth, Botrytis infection |
| Exp. IX | soil batch 5 | square<sup>3</sup> | 2.5 | 2x | 5 | Shoot growth, Botrytis infection |
| Exp. X | soil batch 4 | square<sup>3</sup> | 0.5 | 2x | 6 | Shoot growth, Botrytis infection |
| Exp. XI | Uncond. soil | square<sup>4</sup> |  | 1x | 6 | Botrytis infection |
| Exp. XII | soil batch 6 | square<sup>4</sup> | 1 | - | 5 | Shoot growth, Flowering |

<sup>1</sup> Fertilisation by watering with 1/3 half strength Hoagland solution diluted in tap water.
<sup>2</sup> 5.5 x 5 cm pot size
<sup>3</sup> 8 x 8 x 8.5 cm pot size
<sup>4</sup> 6 x 6 x 6.5 cm pot size

**Table S3.** List of differently abundant bacterial ASVs in soil. Relative abundance and taxonomic information of differently abundant bacterial ASVs (bASVs) from BXplus and BXminus soil. Due to spatial restrictions, kingdom and order are not shown. The phylum/class column reports the phylum infor- mation, but in the case of the Pseudomonadota phylum, the class information is given (Gammaproteobacteria, Alphapro- teobacteria). The mean relative abundances of each bASV in native BXplus and BXminus soil are indicated, ordered by decreasing abundance in the BXplus condition. bASVs with higher abundance in native BXplus soil are shown in bold. C. = candidatus. sg1 = subgroup 1.

| ASV | phylum / class | family | genus | abund.<br>BX+ | abund.<br>BX- |
| --- | --- | --- | --- | --- | --- |
| **bASV143** | Chloroflexota | unassigned | unassigned | 0.69% | 0.24% |
| **bASV136** | Gammaproteobacteria | SC-I-84 | unassigned | 0.59% | 0.35% |
| **bASV105** | Alphaproteobacteria | Sphingomonadaceae | Sphingomonas | 0.47% | 0.24% |
| **bASV319** | Alphaproteobacteria | Sphingomonadaceae | Sphingomonas | 0.30% | 0.18% |
| **bASV231** | Alphaproteobacteria | Sphingomonadaceae | Sphingomonas | 0.27% | 0.14% |
| **bASV343** | Acidobacteriota | unassigned | unassigned | 0.26% | 0.14% |
| **bASV199** | Alphaproteobacteria | Sphingomonadaceae | Sphingomonas | 0.23% | 0.10% |
| **bASV405** | Alphaproteobacteria | Sphingomonadaceae | Sphingomonas | 0.23% | 0.09% |
| **bASV561** | Acidobacteriota | Acidobacteriaceae (sg1) | Occallatibacter | 0.14% | 0.07% |
| **bASV492** | Alphaproteobacteria | Sphingomonadaceae | Sphingomonas | 0.13% | 0.03% |
| **bASV1079** | Gammaproteobacteria | Rhodanobacteraceae | Rhodanobacter | 0.09% | 0.01% |
| **bASV1675** | Gemmatimonadota | Gemmatimonadaceae | unassigned | 0.06% | 0.01% |
| **bASV1830** | Alphaproteobacteria | Sphingomonadaceae | Croceibacterium | 0.05% | 0% |
| **bASV1594** | Bacteroidota | env.OPS 17 | unassigned | 0.03% | 0% |
| **bASV2572** | Acidobacteriota | Solibacteraceae | C. Solibacter | 0.03% | 0% |
| **bASV3104** | Acidobacteriota | Bryobacteraceae | Bryobacter | 0.02% | 0% |
| bASV1032 | Myxococcota | Nannocystaceae | Nannocystis | 0% | 0.03% |
| bASV1914 | Gammaproteobacteria | Rhodocyclaceae | C. Accumulibacter | 0% | 0.05% |
| bASV710 | Actinomycetota | unassigned | unassigned | 0% | 0.09% |
| bASV1356 | Gammaproteobacteria | Rhodocyclaceae | Azovibrio | 0% | 0.08% |
| bASV1762 | Thermodesulfobacteriota | Geobacteraceae | Geomonas | 0% | 0.05% |
| bASV1781 | Gammaproteobacteria | Unknown Family | Acidibacter | 0% | 0.05% |
| bASV1821 | Actinomycetota | Gaiellaceae | Gaiella | 0% | 0.04% |
| bASV3182 | Bacteroidota | Saprospiraceae | unassigned | 0% | 0.02% |

**Table S4.**
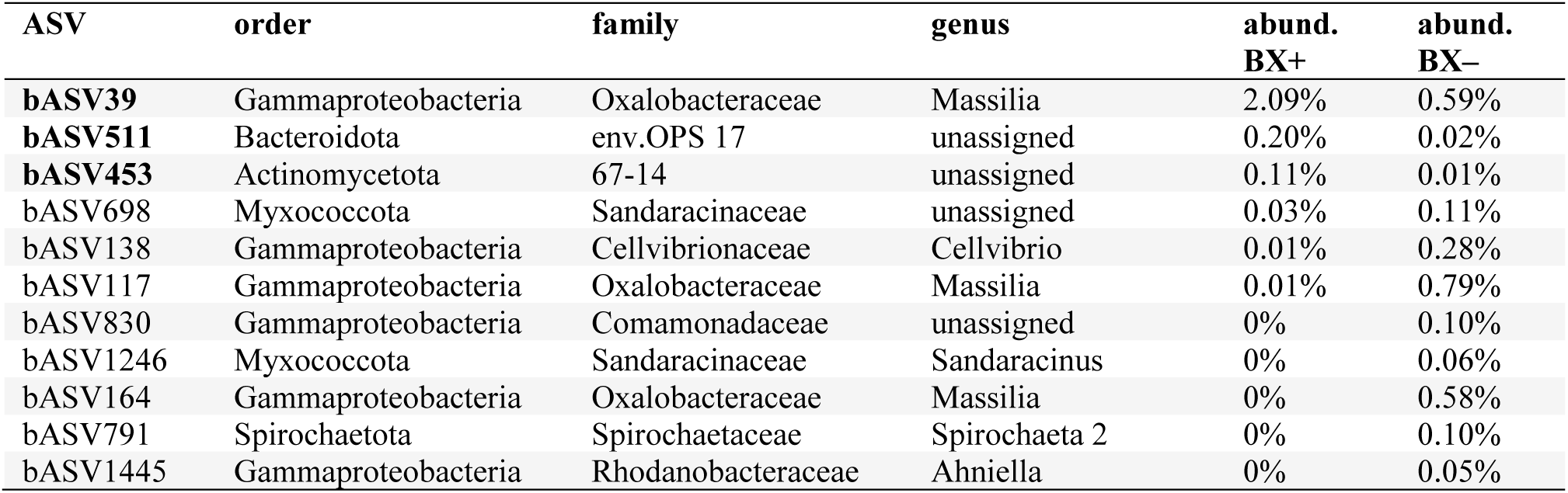
List of differently abundant bacterial ASVs on roots of Arabidopsis. Relative abundance and taxonomic information of differently abundant bacterial ASVs (bASVs) of Arabidopsis root communities from native BXplus and BXminus soil. Due to spatial restrictions, kingdom and order are not shown. The phylum/class column reports the phylum information, but in the case of the Pseudomonadota phylum, the class infor- mation is given (Gammaproteobacteria).The mean relative abundances of each bASV on roots grown in native BXplus and BXminus soil are indicated, ordered by decreasing abundance in the BXplus condition. bASVs with higher abundance on roots from native BXplus soil are shown in bold.

| ASV | order | family | genus | abund.<br>BX+ | abund.<br>BX- |
| --- | --- | --- | --- | --- | --- |
| **bASV39** | Gammaproteobacteria | Oxalobacteraceae | Massilia | 2.09% | 0.59% |
| **bASV511** | Bacteroidota | env.OPS 17 | unassigned | 0.20% | 0.02% |
| **bASV453** | Actinomycetota | 67-14 | unassigned | 0.11% | 0.01% |
| bASV698 | Myxococcota | Sandaracinaceae | unassigned | 0.03% | 0.11% |
| bASV138 | Gammaproteobacteria | Cellvibrionaceae | Cellvibrio | 0.01% | 0.28% |
| bASV117 | Gammaproteobacteria | Oxalobacteraceae | Massilia | 0.01% | 0.79% |
| bASV830 | Gammaproteobacteria | Comamonadaceae | unassigned | 0% | 0.10% |
| bASV1246 | Myxococcota | Sandaracinaceae | Sandaracinus | 0% | 0.06% |
| bASV164 | Gammaproteobacteria | Oxalobacteraceae | Massilia | 0% | 0.58% |
| bASV791 | Spirochaetota | Spirochaetaceae | Spirochaeta 2 | 0% | 0.10% |
| bASV1445 | Gammaproteobacteria | Rhodanobacteraceae | Ahniella | 0% | 0.05% |

The **Supplementary Tables S5 to S8** are in a common excel file with the following work sheets:

**Table S5 Shoot transcriptome** List of significantly differentially expressed genes (*p*adj < 0.05, log2FoldChange ±>1) in shoots as inferred with DESeq2. The work sheet lists the Gene-ID, its base mean of expression, the log2 fold change and its standard error, the DESeq statistic, its *p-*value and adjusted *p-*value.

**Table S6 Root transcriptome** List of significantly differentially expressed genes (*p*adj < 0.05, log2FoldChange ±>1) in roots as inferred with DESeq2. The work sheet lists the Gene-ID, its base mean of expression, the log2 fold change and its standard error, the DESeq statistic, its *p-*value and adjusted *p-*value.

**Table S7 Shoot co-expression analysis** The table lists for shoots for each cluster of co-expressed genes the enriched GO terms with their ID, the description (biological processes), its gene ratio, the background gene ratio, its *p-*value, adjusted *p-*value and Q-value together with the IDs of the identified genes and their count per cluster.

**Table S8 Root co-expression analysis** The table lists for roots for each cluster of co-expressed genes the enriched GO terms with their ID, the description (bio- logical processes), its gene ratio, the background gene ratio, its *p-*value, adjusted *p-*value and Q-value together with the IDs of the identified genes and their count per cluster.

**Table S9.** List of exemplary genes from the RNASeq analysis. List of exemplary genes differentially regulated in shoots and roots of Arabidopsis grown on BXplus compared to BXminus soil. Genes are sorted by tissue (shoots, roots) and by regulation (upregulated in tissue grown on BXplus soil, downregu- lated in tissue grown on BXplus soil). For each gene, the cluster affiliation as well as the log2 fold change (Log2FC) of the gene expression and the adjusted *p-*value for the differential expression on BXplus versus BXminus are indicated.

| Genes in shoots, upregulated (BX<sub>plus</sub> > BX<sub>minus</sub>) |  |  |  |  |
| --- | --- | --- | --- | --- |
| Gene name | Locus | cluster | Log<sub>2</sub>FC | adjusted *p*-value |
| NFYA5 | AT1G54160 | cluster 3 | 1.27 | < 0.0001 |
| AKS1 | AT1G51140 | cluster 3 | 1.02 | < 0.0001 |
| PR5 | AT1G75040 | cluster 2 | 1.28 | 0.03 |
| PR2 | AT3G57260 | cluster 2 | 1.75 | 0.002 |
| Genes in shoots, downregulated (BX<sub>plus</sub> < BX<sub>minus</sub>) |  |  |  |  |
| Gene name | Locus | cluster | Log<sub>2</sub>FC | adjusted *p*-value |
| LHCB6 | AT1G15820 | cluster10 | -1.25 | 0.003 |
| RBCS1A | AT1G67090 | cluster10 | -1.3 | 0.006 |
| CHLI2 | AT5G45930 | cluster10 | -1.04 | 0.003 |
| TN8 | AT1G72910 | cluster 13 | -1.83 | 0.0008 |
| TN9 | AT1G72920 | cluster 13 | -2.60 | < 0.0001 |
| TAA1 | AT1G70560 | cluster 9 | -1.02 | < 0.0001 |
| LAX3 | AT1G77690 | cluster 9 | -1.06 | 0.002 |
| Genes in roots, upregulated (BX<sub>plus</sub> > BX<sub>minus</sub>) |  |  |  |  |
| Gene name | Locus | cluster | Log<sub>2</sub>FC | adjusted *p*-value |
| MYB66/WER | AT5G14750 | cluster 3 | 1.24 | 0.02 |
| MYB23 | AT5G40330 | cluster 3 | 1.47 | 0.002 |
| PRR1 | AT1G32100 | cluster 4 | 1.26 | < 0.0001 |
| MYB43 | AT5G16600 | cluster 4 | 1.10 | 0.0001 |
| SWEET4 | AT3G28007 | cluster 1 | 3.22 | 0.004 |
| SWEET12 | AT5G23660 | cluster 1 | 2.73 | 0.0003 |
| Genes in roots, downregulated (BX<sub>plus</sub> < BX<sub>minus</sub>) |  |  |  |  |
| Gene name | Locus | cluster | Log<sub>2</sub>FC | adjusted *p*-value |
| MKK7 | AT1G18350 | cluster 11 | -4.72 | 0.02 |
| WRKY40 | AT1G80840 | cluster 11 | -2.68 | 0.02 |
| CRK9 | AT4G23170 | cluster 11 | -3.75 | 0.005 |
| TN7 | AT1G72900 | cluster 11 | -2.74 | 0.001 |
| TN8 | AT1G72910 | cluster 11 | -4.31 | < 0.0001 |
| FMO GS-OX1 | AT1G65860 | cluster 10 | -1.71 | 0.03 |
| IMD1 | AT5G14200 | cluster 10 | -2.00 | 0.01 |

### Supplementary Datasets

#### Dataset S1 Microbiota analysis

The Dataset S1 presents the R markdown output of the microbiota analysis. The raw sequencing data is available from ENA, the input data, R code and R markdown output are available from GitHub under https://github.com/PMI-Basel/Stengele_et_al_At_BX-feedbacks.

#### Dataset S2 Transcriptome analysis

The Dataset S2 presents the R markdown output of the transcriptome analysis. The raw sequencing data is available from ENA, input data, R code and R markdown output are available from GitHub under https://github.com/PMI-Basel/Stengele_et_al_At_BX-feedbacks.

## References

1. Badri DV, Zolla G, Bakker MG, Manter DK, Vivanco JM. 2013. Potential impact of soil microbiomes on the leaf metabolome and on herbivore feeding behavior. New Phytologist 198: 264–273.

2. Bai B, Liu W, Qiu X, Zhang J, Zhang J, Bai Y. 2022. The root microbiome: Community assembly and its contributions to plant fitness. Journal of Integrative Plant Biology 64: 230–243.

3. Beilsmith K, Perisin M, Bergelson J. 2021. Natural Bacterial Assemblages in Arabidopsis thaliana Tissues Become More Distinguishable and Diverse during Host Development. mBio 12: e02723–20.

4. Berns AE, Philipp H, Narres H-D, Burauel P, Vereecken H, Tappe W. 2008. Effect of gamma-sterilization and autoclaving on soil organic matter structure as studied by solid state NMR, UV and fluorescence spectroscopy. European Journal of Soil Science 59: 540–550.

5. Bezemer TM, Lawson CS, Hedlund K, Edwards AR, Brook AJ, Igual JM, Mortimer SR, Van Der Putten WH. 2006. Plant species and functional group effects on abiotic and microbial soil properties and plant–soil feedback responses in two grasslands. Journal of Ecology 94: 893–904.

6. Bray JR, Curtis JT. 1957. An ordination of the upland forest communities of southern wisconsin. Ecological Monographs 27: 325–349.

7. Bulgarelli D, Schlaeppi K, Spaepen S, van Themaat EVL, Schulze-Lefert P. 2013. Structure and functions of the bacterial microbiota of plants. Annual Review of Plant Biology 64: 807–838.

8. Cadot S, Gfeller V, Hu L, Singh N, Sánchez-Vallet A, Glauser G, Croll D, Erb M, Heijden MGA, Schlaeppi K. 2021. Soil composition and plant genotype determine benzoxazinoid-mediated plant–soil feedbacks in cereals. Plant, Cell & Environment 44: 3732–3744.

9. Cardoso C, Charnikhova T, Jamil M, Delaux P-M, Verstappen F, Amini M, Lauressergues D, Ruyter- Spira C, Bouwmeester H. 2014. Differential activity of Striga hermonthica seed germination stimulants and Gigaspora rosea hyphal branching factors in rice and their contribution to underground communication (M Öpik, Ed.). PLoS ONE 9: e104201.

10. Cotton TEA, Pétriacq P, Cameron DD, Meselmani MA, Schwarzenbacher R, Rolfe SA, Ton J. 2019. Metabolic regulation of the maize rhizobiome by benzoxazinoids. The ISME Journal 13: 1647–1658.

11. Delory BM, Callaway RM, Semchenko M. 2024. A trait-based framework linking the soil metabolome to plant–soil feedbacks. New Phytologist 241: 1910–1921.

12. Dobin A, Davis CA, Schlesinger F, Drenkow J, Zaleski C, Jha S, Batut P, Chaisson M, Gingeras TR. 2013. STAR: ultrafast universal RNA-seq aligner. Bioinformatics 29: 15–21.

13. Durán P, Thiergart T, Garrido-Oter R, Agler M, Kemen E, Schulze-Lefert P, Hacquard S. 2018. Microbial interkingdom interactions in roots promote Arabidopsis survival. Cell 175: 973–983.e14.

14. Fernandes AD, Macklaim JM, Linn TG, Reid G, Gloor GB. 2013. ANOVA-like differential expression (ALDEx) analysis for mixed population RNA-Seq. PLOS ONE 8: e67019.

15. Fernandes AD, Reid JN, Macklaim JM, McMurrough TA, Edgell DR, Gloor GB. 2014. Unifying the analysis of high-throughput sequencing datasets: characterizing RNA-seq, 16S rRNA gene sequencing and selective growth experiments by compositional data analysis. Microbiome 2: 15.

16. French E, Kaplan I, Iyer-Pascuzzi A, Nakatsu CH, Enders L. 2021. Emerging strategies for precision microbiome management in diverse agroecosystems. Nature Plants 7: 256–267.

17. Gfeller V, Cadot S, Waelchli J, Gulliver S, Terrettaz C, Thönen L, Mateo P, Robert CAM, Mascher F, Steinger T, et al. 2023a. Soil chemical and microbial gradients determine accumulation of root-exuded secondary metabolites and plant–soil feedbacks in the field. Journal of Sustainable Agriculture and Environment: sae2.12063.

18. Gfeller V, Waelchli J, Pfister S, Deslandes-Hérold G, Mascher F, Glauser G, Aeby Y, Mestrot A, Robert CA, Schlaeppi K, et al. 2023b. Plant secondary metabolite-dependent plant-soil feedbacks can improve crop yield in the field. eLife 12: e84988.

19. Gimsing AL, Kirkegaard JA. 2009. Glucosinolates and biofumigation: fate of glucosinolates and their hydrolysis products in soil. Phytochemistry Reviews 8: 299–310.

20. Goossens P, Spooren J, Baremans KCM, Andel A, Lapin D, Echobardo N, Pieterse CMJ, Van Den Ackerveken G, Berendsen RL. 2023. Obligate biotroph downy mildew consistently induces near-identical protective microbiomes in Arabidopsis thaliana. Nature Microbiology 8: 2349–2364.

21. Govrin EM, Levine A. 2002. Infection of Arabidopsis with a necrotrophic pathogen, Botrytis cinerea, elicits various defense responses but does not induce systemic acquired resistance (SAR). Plant Molecular Biology 48: 267–276.

22. Hannula SE, Heinen R, Huberty M, Steinauer K, De Long JR, Jongen R, Bezemer TM. 2021. Persistence of plant-mediated microbial soil legacy effects in soil and inside roots. Nature Communications 12: 5686.

23. Harris CJ, Amtmann A, Ton J. 2023. Epigenetic processes in plant stress priming: Open questions and new approaches. Current Opinion in Plant Biology 75: 102432.

24. Hu L, Robert CAM, Cadot S, Zhang X, Ye M, Li B, Manzo D, Chervet N, Steinger T, van der Heijden MGA, et al. 2018. Root exudate metabolites drive plant-soil feedbacks on growth and defense by shaping the rhizosphere microbiota. Nature Communications 9: 2738.

25. Hubbard CJ, Li B, McMinn R, Brock MT, Maignien L, Ewers BE, Kliebenstein D, Weinig C. 2019. The effect of rhizosphere microbes outweighs host plant genetics in reducing insect herbivory. Molecular Ecology 28: 1801–1811.

26. Hüther P, Schandry N, Jandrasits K, Bezrukov I, Becker C. 2020. ARADEEPOPSIS, an automated workflow for top-view plant phenomics using semantic segmentation of leaf states. The Plant Cell 32: 3674– 3688.

27. Inzé D, Nelissen H. 2022. The translatability of genetic networks from model to crop species: lessons from the past and perspectives for the future. New Phytologist 236: 43–48.

28. Jacoby RP, Koprivova A, Kopriva S. 2021. Pinpointing secondary metabolites that shape the composition and function of the plant microbiome. Journal of Experimental Botany 72: 57–69.

29. Janse Van Rensburg H, Schandry N, Waelchli J, Stengele K, Cadot S, Jandrasits K, Becker C, Schlaeppi K. 2025. A TNL receptor mediates microbiome feedbacks in Arabidopsis.

30. Janse van Rensburg H, Stengele K, Schlaeppi K. 2024. Understanding plant responsiveness to microbiome feedbacks. Current Opinion in Plant Biology 81: 102603.

31. Jing J, Cong W-F, Bezemer TM. 2022. Legacies at work: plant–soil–microbiome interactions underpinning agricultural sustainability. Trends in Plant Science 27: 781–792.

32. Karasov TL, Chae E, Herman JJ, Bergelson J. 2017. Mechanisms to mitigate the trade-off between growth and defense. The Plant Cell 29: 666–680.

33. Korenblum E, Massalha H, Aharoni A. 2022. Plant–microbe interactions in the rhizosphere via a circular metabolic economy. The Plant Cell 34: 3168–3182.

34. Kudjordjie EN, Sapkota R, Steffensen SK, Fomsgaard IS, Nicolaisen M. 2019. Maize synthesized benzoxazinoids affect the host associated microbiome. Microbiome 7: 59.

35. Li Z, Wang Z, Zhang Y, Yang J, Guan K, Song Y. 2025. Identification of stress-alleviating strains from the core drought-responsive microbiome of Arabidopsis ecotypes. The ISME Journal 19: wraf067.

36. Lin H, Peddada SD. 2020. Analysis of compositions of microbiomes with bias correction. Nature Communications 11: 3514.

37. van Loon LC, Rep M, Pieterse CMJ. 2006. Significance of inducible defense-related proteins in infected plants. Annual Review of Phytopathology 44: 135–162.

38. Love MI, Huber W, Anders S. 2014. Moderated estimation of fold change and dispersion for RNA-seq data with DESeq2. Genome biology 15: 1–21.

39. Maag D, Köhler A, Robert CAM, Frey M, Wolfender J-L, Turlings TCJ, Glauser G, Erb M. 2016. Highly localized and persistent induction of Bx1-dependent herbivore resistance factors in maize. The Plant Journal 88: 976–991.

40. Mallick H, Rahnavard A, McIver LJ, Ma S, Zhang Y, Nguyen LH, Tickle TL, Weingart G, Ren B, Schwager EH, et al. 2021. Multivariable association discovery in population-scale meta-omics studies. PLoS computational biology 17: e1009442.

41. Mariotte P, Mehrabi Z, Bezemer TM, De Deyn GB, Kulmatiski A, Drigo B, Veen GF (Ciska), van der Heijden MGA, Kardol P. 2018. Plant–soil feedback: Bridging natural and agricultural sciences. Trends in Ecology & Evolution 33: 129–142.

42. Martinez-Medina A, Flors V, Heil M, Mauch-Mani B, Pieterse CMJ, Pozo MJ, Ton J, van Dam NM, Conrath U. 2016. Recognizing plant defense priming. Trends in Plant Science 21: 818–822.

43. McLaughlin S, Joller C, Siffert A, Stirnemann EM, Sasse J. 2023. A Versatile Glass Jar System for Semihydroponic Root Exudate Profiling. Journal of Visualized Experiments: 66070.

44. McMurdie PJ, Holmes S. 2013. phyloseq: An R package for reproducible interactive analysis and graphics of microbiome census data. PLOS ONE 8: e61217.

45. Niculaes C, Abramov A, Hannemann L, Frey M. 2018. Plant protection by Benzoxazinoids—recent insights into biosynthesis and function. Agronomy 8: 143.

46. Niemeyer HM. 2009. Hydroxamic acids derived from 2-Hydroxy-2 H -1,4-Benzoxazin-3(4 H)-one: Key defense chemicals of cereals. Journal of Agricultural and Food Chemistry 57: 1677–1696.

47. Paulson JN, Stine OC, Bravo HC, Pop M. 2013. Differential abundance analysis for microbial marker-gene surveys. Nature Methods 10: 1200–1202.

48. Pieterse CMJ, Van Wees SCM, Hoffland, Ellis, Van Pelt JA, Loon LC van. 1996. Systemic resistance in Arabidopsis induced by biocontrol bacteria is independent of salicylic acid accumulation and pathogenesis- related gene expression. The Plant Cell 8: 1225–1237.

49. Pieterse CM, van Wees SC, van Pelt JA, Knoester M, Laan R, Gerrits H, Weisbeek PJ, van Loon LC. 1998. A novel signaling pathway controlling induced systemic resistance in Arabidopsis. The Plant Cell 10: 1571–1580.

50. Pieterse CMJ, Zamioudis C, Berendsen RL, Weller DM, Van Wees SCM, Bakker PAHM. 2014. Induced systemic resistance by beneficial microbes. Annual Review of Phytopathology 52: 347–375.

51. Pineda A, Kaplan I, Hannula SE, Ghanem W, Bezemer TM. 2020. Conditioning the soil microbiome through plant–soil feedbacks suppresses an aboveground insect pest. New Phytologist 226: 595–608.

52. Posit team. 2023. RStudio: Integrated Development Environment for R. Boston, MA: Posit Software, PBC.

53. van der Putten WH, Bardgett RD, Bever JD, Bezemer TM, Casper BB, Fukami T, Kardol P, Klironomos JN, Kulmatiski A, Schweitzer JA, et al. 2013. Plant–soil feedbacks: the past, the present and future challenges. Journal of Ecology 101: 265–276.

54. R Core Team. 2017. R: A language and environment for statistical computing. Vienna, Austria: R Foundation for Statistical Computing.

55. Rau A, Maugis-Rabusseau C. 2018. Transformation and model choice for RNA-seq co-expression analysis. Briefings in bioinformatics 19: 425–436.

56. Robert CAM, Mateo P. 2022. The chemical ecology of Benzoxazinoids. CHIMIA 76: 928.

57. Rodriguez RE, Mecchia MA, Debernardi JM, Schommer C, Weigel D, Palatnik JF. 2010. Control of cell proliferation in Arabidopsis thaliana by microRNA miR396. Development (Cambridge, England) 137: 103–112.

58. Ross AF. 1961. Systemic acquired resistance induced by localized virus infections in plants. Virology 14: 340– 358.

59. Ruijter JM, Ramakers C, Hoogaars WMH, Karlen Y, Bakker O, van den Hoff MJB, Moorman AFM. 2009. Amplification efficiency: linking baseline and bias in the analysis of quantitative PCR data. Nucleic Acids Research 37: e45.

60. Ryals JA, Neuenschwander UH, Willits MG, Molina A, Steiner H-Y, Hunt MD. 1996. Systemic acquired resistance. The Plant Cell 8: 1809–1819.

61. Salas-González I, Reyt G, Flis P, Custódio V, Gopaulchan D, Bakhoum N, Dew TP, Suresh K, Franke RB, Dangl JL, et al. 2021. Coordination between microbiota and root endodermis supports plant mineral nutrient homeostasis. Science 371: eabd0695.

62. Santos LF, Olivares FL. 2021. Plant microbiome structure and benefits for sustainable agriculture. Current Plant Biology 26: 100198.

63. Sasse J, Martinoia E, Northen T. 2018. Feed your friends: Do plant exudates shape the root microbiome? Trends in Plant Science 23: 25–41.

64. Schandry N, Becker C. 2020. Allelopathic plants: Models for studying plant–interkingdom interactions. Trends in Plant Science 25: 176–185.

65. Schindelin J, Arganda-Carreras I, Frise E, Kaynig V, Longair M, Pietzsch T, Preibisch S, Rueden C, Saalfeld S, Schmid B, et al. 2012. Fiji: an open-source platform for biological-image analysis. Nature Methods 9: 676–682.

66. Schmitz L, Yan Z, Schneijderberg M, de Roij M, Pijnenburg R, Zheng Q, Franken C, Dechesne A, Trindade LM, van Velzen R, et al. 2022. Synthetic bacterial community derived from a desert rhizosphere confers salt stress resilience to tomato in the presence of a soil microbiome. The ISME Journal 16: 1907–1920.

67. Schneider CA, Rasband WS, Eliceiri KW. 2012. NIH Image to ImageJ: 25 years of image analysis. Nature Methods 9: 671–675.

68. Semchenko M, Barry KE, Vries FT, Mommer L, Moora M, Maciá-Vicente JG. 2022. Deciphering the role of specialist and generalist plant–microbial interactions as drivers of plant–soil feedback. New Phytologist 234: 1929–1944.

69. Shigenaga AM, Argueso CT. 2016. No hormone to rule them all: Interactions of plant hormones during the responses of plants to pathogens. Seminars in Cell & Developmental Biology 56: 174–189.

70. Spooren J, Van Bentum S, Thomashow LS, Pieterse CMJ, Weller DM, Berendsen RL. 2024. Plant-driven assembly of disease-suppressive soil microbiomes. Annual Review of Phytopathology 62: 1–30.

71. Stringlis IA, Yu K, Feussner K, de Jonge R, Van Bentum S, Van Verk MC, Berendsen RL, Bakker PAHM, Feussner I, Pieterse CMJ. 2018. MYB72-dependent coumarin exudation shapes root microbiome assembly to promote plant health. Proceedings of the National Academy of Sciences 115.

72. Teasdale JR, Rice CP, Cai G, Mangum RW. 2012. Expression of allelopathy in the soil environment: soil concentration and activity of benzoxazinoid compounds released by rye cover crop residue. Plant Ecology 213: 1893–1905.

73. Trivedi P, Leach JE, Tringe SG, Sa T, Singh BK. 2020. Plant–microbiome interactions: from community assembly to plant health. Nature Reviews Microbiology 18: 607–621.

74. Venturelli S, Belz RG, Kämper A, Berger A, Von Horn K, Wegner A, Böcker A, Zabulon G, Langenecker T, Kohlbacher O, et al. 2015. Plants release precursors of histone deacetylase inhibitors to suppress growth of competitors. The Plant Cell 27: 3175–3189.

75. Wang D, He X, Baer M, Lami K, Yu B, Tassinari A, Salvi S, Schaaf G, Hochholdinger F, Yu P. 2024. Lateral root enriched Massilia associated with plant flowering in maize. Microbiome 12: 124.

76. Weiss SJ, Xu Z, Amir A, Peddada S, Bittinger K, Gonzalez A, Lozupone C, Zaneveld JR, Vazquez-Baeza Y, Birmingham A, et al. 2015. Effects of library size variance, sparsity, and compositionality on the analysis of microbiome data.

77. Wouters FC, Blanchette B, Gershenzon J, Vassão DG. 2016. Plant defense and herbivore counter-defense: benzoxazinoids and insect herbivores. Phytochemistry Reviews 15: 1127–1151.

78. Wu T, Hu E, Xu S, Chen M, Guo P, Dai Z, Feng T, Zhou L, Tang W, Zhan L, et al. 2021. clusterProfiler 4.0: A universal enrichment tool for interpreting omics data. Innovation (Cambridge (Mass.)) 2: 100141.

79. Yu P, He X, Baer M, Beirinckx S, Tian T, Moya YAT, Zhang X, Deichmann M, Frey FP, Bresgen V, et al. 2021. Plant flavones enrich rhizosphere Oxalobacteraceae to improve maize performance under nitrogen deprivation. Nature Plants 7: 481–499.

80. Zeier J. 2021. Metabolic regulation of systemic acquired resistance. Current Opinion in Plant Biology 62: 102050.

## Supplementary References

Abarenkov K, Nilsson RH, Larsson K-H, Taylor AFS, May TW, Frøslev TG, Pawlowska J, Lindahl B, Põldmaa K, Truong C, et al. 2024. The UNITE database for molecular identification and taxonomic communication of fungi and other eukaryotes: sequences, taxa and classifications reconsidered. Nucleic Acids Research 52: D791–D797.

Callahan BJ, McMurdie PJ, Rosen MJ, Han AW, Johnson AJA, Holmes SP. 2016. DADA2: High-resolution sam- ple inference from Illumina amplicon data. Nature Methods 13: 581–583.

Lindsey III BE, Rivero L, Calhoun CS, Grotewold E, Brkljacic J. 2017. Standardized method for high-throughput sterilization of Arabidopsis seeds. Journal of Visualized Experiments: 56587.

Martin M. 2011. Cutadapt removes adapter sequences from high-throughput sequencing reads. EMBnet.journal 17: 10–12.

McLaughlin S, Joller C, Siffert A, Stirnemann EM, Sasse J. 2023. A Versatile Glass Jar System for Semihydroponic Root Exudate Profiling. Journal of Visualized Experiments: 66070.

Quast C, Pruesse E, Yilmaz P, Gerken J, Schweer T, Yarza P, Peplies J, Glöckner FO. 2013. The SILVA riboso- mal RNA gene database project: improved data processing and web-based tools. Nucleic Acids Research 41: D590– D596.

R Core Team. 2017. R: A language and environment for statistical computing. Vienna, Austria: R Foundation for Statistical Computing.

